# Inverse regulation of secretion and inflammation in human airway gland serous cells by neuropeptides upregulated in allergy and asthma

**DOI:** 10.1101/632224

**Authors:** Derek B. McMahon, Michael A. Kohanski, Charles C.L. Tong, Peter Papagiannopoulos, Nithin D. Adappa, James N. Palmer, Robert J. Lee

## Abstract

Airway submucosal gland serous cells are sites of expression of the cystic fibrosis transmembrane conductance regulator (CFTR) and are important for fluid secretion in conducting airways from the nose down to small bronchi. We tested if serous cells from human nasal turbinate glands secrete bicarbonate (HCO_3_^−^), important for mucus polymerization, during stimulation with the cAMP-elevating agonist vasoactive intestinal peptide (VIP) and if this requires CFTR. Isoalted serous cells stimulated with VIP exhibited a ~20% cAMP-dependent decrease in cell volume and a ~0.15 unit decrease in intracellular pH (pH_i_), reflecting activation of Cl^−^ and HCO_3_^−^ secretion, respectively. Pharmacology, ion substitution, and studies using cells from CF patients suggest serous cell HCO_3_^−^ secretion is mediated by conductive efflux directly through CFTR. Interestingly, we found that neuropeptide Y (NPY) reduced VIP-evoked secretion by blunting cAMP increases and reducing CFTR activation through G_i_-coupled NPY1R. Culture of primary gland serous cells in a model that maintained a serous phenotype confirmed the activating and inhibiting effects of VIP and NPY, respectively, on fluid and HCO_3_^−^ secretion. Moreover, VIP enhanced secretion of antimicrobial peptides and antimicrobial efficacy of gland secretions while NPY reduced antimicrobial secretions. In contrast, NPY enhanced the release of cytokines during inflammatory stimuli while VIP reduced cytokine release through a mechanism requiring CFTR conductance. As levels of VIP and NPY are up-regulated in disease like allergy, asthma, and chronic rhinosinusitis, the balance of these two peptides in the airway may control airway mucus rheology and inflammatory responses through gland serous cells.

## INTRODUCTION

Several distinct obstructive airway diseases share a phenotype of thickened mucus and/or mucostasis, including chronic rhinosinusitis (CRS) (1), cystic fibrosis (CF), asthma (2, 3), and COPD (4). In conducting airways from the nasal turbinates down to small bronchi ~1 mm^2^ in diameter, a large percentage of airway surface liquid (ASL) and mucus is generated in airway submucosal exocrine glands (5-7). Submucosal gland serous acinar cells are sites of expression of the cystic fibrosis (CF) transmembrane conductance regulator (CFTR) Cl^−^ channel (8-12). Defects in CFTR-dependent serous cell secretion likely play an important role in CF pathology, supported by observations of occluded mucus-filled gland ducts, gland hypertrophy and hyperplasia, and gland infection in lungs of CF patients (13, 14). Intact glands from CF individuals or transgenic CF animals secrete less fluid in response to cAMP-elevating agonists such as vasoactive intestinal peptide (VIP) (15-24) compared with non-CF glands. Gland hypertrophy, duct plugging, and/or excess mucus secretion have also been observed in COPD and asthma (4, 25-35), with gland hypertrophy being greater in fatal asthma cases than non-fatal cases (33).

Proper gland secretion likely requires bicarbonate (HCO_3_^−^) secretion by serous cells at the distal ends of the glands to facilitate polymerization of mucins secreted by more proximal mucous cells (36-41) (**Figure 1A**). However, the mechanisms by which serous cells secrete HCO_3_^−^ are unknown. HCO_3_^−^ may also be critical to the efficacy of antimicrobial peptides secreted by serous cells (42-45), including lysozyme, lactoferrin, LL-37, and Muc7 (46). Understanding how airway glands secrete HCO_3_^−^ may yield insights into the pathophysiology of CRS, CF, COPD, and asthma, all of which have a common phenotype of altered airway mucus secretion or rheology.

**Figure 1:**
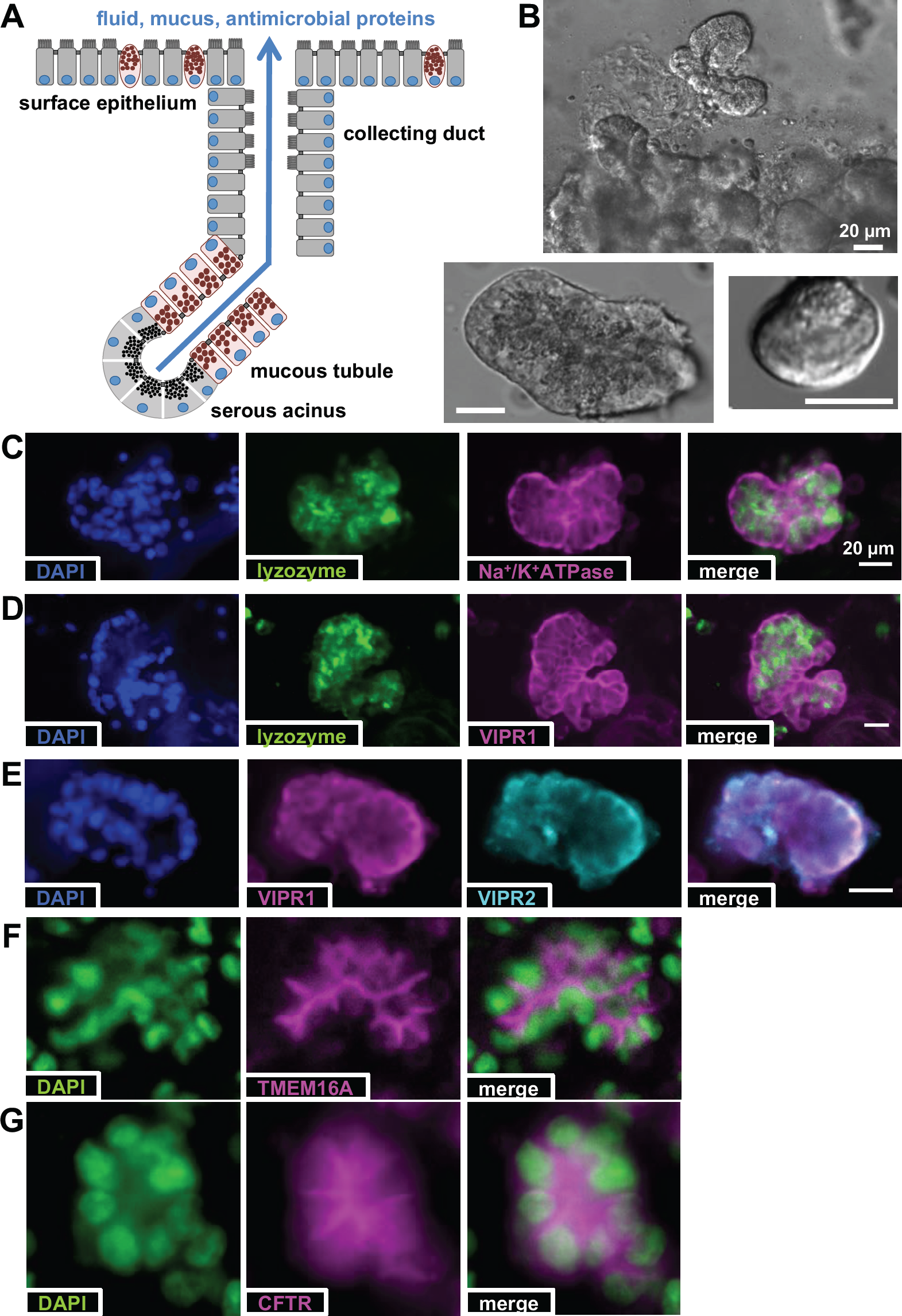
Isolated airway serous acinar cells. **(A).** Representative diagram showing serous acinar cells at the distal ends of submucosal glands, which secrete the bulk of fluid in response to agonists that utilize cAMP or Ca^2+^ as second messengers. **(B).** Primary human serous acini and acinar cells isolated from human middle turbinate samples. **(C-E).** Isolated serous acini exhibited punctate granular immunofluorescence for lysozyme as well as basolateral membrane staining for Na^+^/K^+^ ATPase, VIPR1, and VIPR2. **(F-G).** Apical membrane staining was observed for secretory Cl-channels TMEM16A and CFTR, as previously described. Scale bars are 20 μm.

We previously developed live cell imaging techniques to study living primary mouse nasal serous cells and demonstrated that they secrete HCO_3_^−^ during cholinergic stimulation (47). Cholinergic-induced secretion is largely intact in CF (10-12, 15-24, 46-48), as it is mediated by Ca^2+^ activated Cl^−^ channels, including TMEM16A (12). An initial goal of the current study was to directly test if serous acinar cells secrete HCO_3_^−^ during stimulation with VIP, whether this occurs through CFTR, and if activation of TMEM16A could substitute. A further goal was to understand the potential relationship of VIP and neuropeptide Y (NPY), both upregulated in inflammatory airway diseases, in control and composition of airway gland secretions. A recent review highlighted a need for a clearer portrait of neuropeptide regulation of submucosal gland secretion within the context of the diverse lung diseases characterized by mucus obstruction (49).

Parasympathetic VIPergic neurons (50-55) and NPY-containing fibers (56-58) are exist in the respiratory tract, with some nerves co-expressing VIP and NPY (59), including in the proximity of submucosal glands (60, 61). Immune cells like activated macrophages (62-64) or epithelial cells (65) can also make NPY. Elevated NPY in allergic asthma (66, 67) may link psychological stress with asthma exacerbations (68-70). Both VIP-containing and NPY-containing nerves may be increased in mucosa from patients with allergic rhinitis (71, 72) or irritative toxic rhinitis (73). VIP and NPY, but not substance P or calcitonin gene-related peptide (CGRP), are found in the pedicle of nasal polyps, suggesting they may play a role in polyp formation (74). Mice lacking NPY or NPY1R have reduced allergic airway inflammation (75), suggesting this neuropeptide and this receptor isoform detrimentally contribute to inflammatory airway diseases. One study found NPY and NPYR1 expression elevated in mouse lungs after influenza infection; knockout of NPY reduced the severity of disease and lowered IL-6 levels (63). In other studies outside the airway, NPY deficiency can reduce Th2 responses (76, 77).

The role of VIP as a cAMP-dependent activator of gland secretion has been extensively studied (11, 19, 78), but the role of NPY is less clear. A cocktail of NPY and norepinephrine inhibited cultured tracheal gland cell glycoprotein secretion (79), and NPY inhibits bulk mucus secretion in ferret trachea (80), though there is little mechanistic data for how NPY affects epithelial or gland cells specifically. We sought to understand how VIP and NPY signaling may interact to control of gland serous cell secretion. Because NPY receptors are often G_i_-coupled, they may reduce cAMP-evoked responses to G_s_-coupled VIP or beta-adrenergic receptors (81-83) or CCK receptors (84). We hypothesized that NPY may reduce airway serous cell fluid and/or HCO_3_^−^ secretion during VIPergic stimulation though modulation of cAMP and thus CFTR. Moreover, VIP and NPY are potent immunomodulators in the gut (85). These peptides may be relevant for airway gland-cell-driven inflammation which may help drive airway submucosal remodeling or airway inflammation.

We examined the effects of VIP and NPY on secretion from primary human airway gland serous acinar cells isolated from nasal turbinate. Cells were studied acutely as well as in an air-liquid interface (ALI) culture model that retained expression of important serous cell markers and facilitated polarized studies and co-culture with human immune cells. Results below contribute to our understanding of airway serous cell secretion and the role of CFTR in both secretion and inflammation, also suggesting therapeutic strategies (NPY1R antagonists, TMEM16A activators) for obstructive inflammatory airway diseases.

## RESULTS

### VIP stimulates both Cl^−^ and HCO_3_^−^ secretion from airway gland serous cells through CFTR

Submucosal gland acini and single acinar cells (**Figure 1B**) were isolated from human nasal middle turbinate as previously described (11). Serous acini exhibited secretory-granule localized immunofluorescence for serous cell marker lysozyme (**Figure 1C**; as previously reported (10, 12, 48)) as well as basolateral immunofluorescence of VIP receptors VIPR1 (VPAC1; **Figure 1D**) and VIPR2 (VPAC2; **Figure 1E**). In contrast, secretory Cl^−^ channels TMEM16A and CFTR exhibited apical membrane immunofluorescence (**Figure 1F-G**), as previously observed (10, 12, 48).

Fluid and ion transport pathways were studied in acutely isolated serous cells using simultaneous DIC measurement of cell volume and quantitative fluorescence microscopy of ion indicator dyes to measure the concentrations of ions involved in driving fluid secretion (Cl^−^/ HCO_3_^−^), a technique pioneered in the study of parotid gland secretory acinar cells (86, 87) and adapted previously for airway gland serous cells (10-12, 46-48). Epithelial fluid secretion is driven largely by Cl^−^. Acinar cell shrinkage during agonist stimulation reflects efflux of cellular K^+^ and Cl^−^ upon activation of secretion and movement of osmotically obliged water. Cell swelling upon removal of agonist reflects solute uptake via mechanisms that sustain secretion such as the bumetanide-sensitive Na^+^K^+^2Cl^−^ co-transporter NKCC1 (46) (**Figure 2A**). Human nasal serous cells shrank by approximately 20% when stimulated with the cAMP-elevating agonists forskolin or VIP (**Figure 2B**), as we previously reported (11). We now found this was also accompanied by a transient decrease in intracellular pH (pH_i_) followed by a more sustained increase in pH_i_ (**Figure 2C-D**).

**Figure 2:**
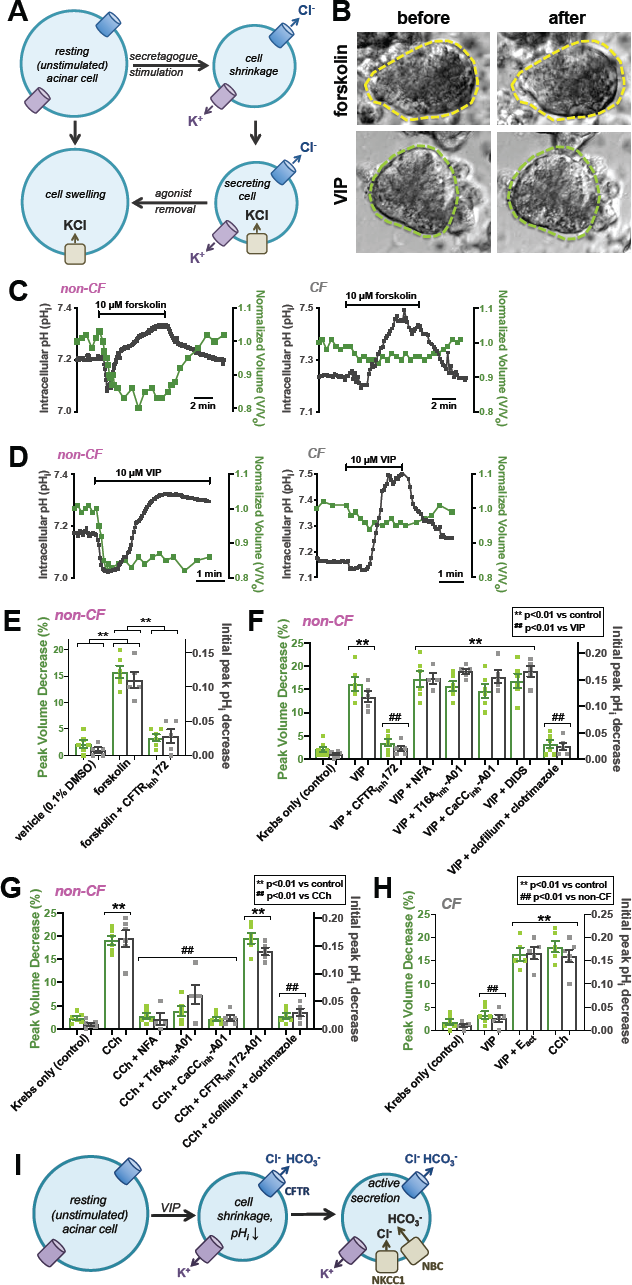
cAMP agonist stimulation of human nasal serous cells results in CFTR-dependent Cl^−^ secretion, revealed by cell shrinkage, concomitant with a CFTR-dependent decrease in pH_i_. **(A)** Diagram showing use of acinar cell volume measurements to track fluid secretion, primarily driven by Cl^−^ secretion, which was combined with simultaneous measurement of pH_i_ to track HCO_3_^−^ secretion. **(B)** Non-CF serous cells stimulated with adenylyl cyclase-activating forskolin (top) or G_s_-coupled receptor agonist VIP (bottom) exhibited ∼15% shrinkage reflecting the activation of fluid secretion. **(C-D)** In cells from non-CF patients, forskolin-induced (*C*) or VIP-induced (*0*) shrinkage (∼15%; green) was accompanied by a transient decrease in pH_i_ (∼0.1 unit; gray) followed by a sustained alkalinization. CF cells exhibited markedly reduced shrinkage and pH_i_ decrease; subsequent alkalinization was intact. **(E-H).** Bar graphs showing peak shrinkage (green) and pH_i_ decrease (gray) in non-CF (*E-G*) and CF (*H*) cells. Forskolin-induced shrinkage was inhibited by CFTR_inh_172 (*E*), while VIP-induced shrinkage was inhibited by CFTR_inh_172 and K^+^ channel inhibitors clofilium and clotrimazole (*F*). Ca^2+^-activated Cl^−^ channel inhibitors NFA, T16A_inh_-A01, CaCC_inh_-A01 or DIDS had no effect on VIP-induced shrinkage (*F*) but did block shrinkage during stimulation with a Ca^2+^-elevating agonist carbachol (CCh). CF cells exhibited minimal responses to VIP but intact response to CCh. VIP responses were restored by TMEM16A-activator E_act_. All experiments done at 37°C with 5% CO_2_/25 mM HCO_3_^−^. All data in *E-H* are mean ± SEM of 5-8 individual experiments from at least 4 individual patients. Significances determined by one-way ANOVA, Bonferroni posttest. **(I)** Diagram showing activation of airway serous cell secretion by VIP, with Cl^−^ and HCO_3_^−^ efflux through CFTR (apically localized in intact glands) causing a decrease in cell volume and pH_i_. Influx of Cl^−^ though NKCC1 and influx of HCO_3_^−^ through NBC (both basolaterally localized in intact glands) maintains the driving force for Cl^−^ and HCO_3_^−^ efflux during sustained secretion.

Both the cell shrinkage and decrease in pH_i_ were absent in cells isolated from CF patients (**Figure 2C-D**). The agonist-evoked pH_i_ decrease was absent when HCO_3_^−^ was removed from the media (**Supplemental Figure 1A-C**), and the secondary pH_i_ increase was blocked with inhibition of the Na^+^HCO_3_^−^ co-transporter (NBC; **Supplemental Figure 1D**). This suggests the transient pH_i_ decrease reflects HCO_3_^−^ efflux during activation of secretion, while the pH_i_ increase reflects activation of NBC, sustaining HCO_3_^−^ secretion by keeping intracellular HCO_3_^−^ high. This is similar to cholinergic evoked pH_i_ decreases and subsequent elevation of pH_i_ by Na^+^/H^+^ exchangers (NHEs) in mouse nasal serous cells (47), but reveals an important mechanistic difference between cAMP and Ca^2+^ pathways.

The pH_i_ decrease was also blocked by eliminating the driving forces for HCO_3_^−^ efflux using ion substitution (**Supplemental Figure 1E**), suggesting the pH_i_ decrease is mediated by conductive HCO_3_^−^ efflux, such as an ion channel. Both forskolin-induced pH_i_ decrease and cell volume decrease were inhibited by CFTR_inh_172 (10 μM; **Figure 2E**). VIP-induced cell volume and pH_i_ decreases were blocked by CFTR_inh_172 or K^+^ channel inhibitors clofilium and clotrimazole (30 μM each; **Figure 2F**) demonstrating a requirement for both CFTR and conductive counterion K^+^ efflux, supporting a Cl^−^ channel as the HCO_3_ ^−^ efflux pathway. VIP-evoked responses were not blocked by the calcium-activated Cl^−^ channel inhibitors niflumic acid (100 μM), T16A_inh_-A01 (10 μM), CaCC_inh_-A01 (10 μM) or 4,4’-Diisothiocyanostilbene-2-2”-disulfonic acid; (DIDS; 1 mM) (**Figure 2F**). These data suggest that VIP receptor activation or direct cAMP elevation with forskolin can activate both Cl^−^ and HCO_3_^−^ secretion directly through CFTR. We found no evidence for Cl^−^/ HCO_3_^−^ exchanger-mediated HCO_3_^−^ efflux in primary serous cells (**Supplemental Figure 2**), suggesting CFTR is the main HCO_3_^−^ efflux pathway during cAMP stimulation, agreeing with recent Calu-3 studies suggesting CFTR sustains HCO_3_^−^secretion (88, 89) instead of the pendrin Cl^−^/ HCO_3_^−^ exchanger (90, 91).

In contrast, cholinergic agonist carbachol (CCh; 10 μM), which activates Ca^2+^-driven TMEM16A-mediated secretion (10-12, 46), stimulated cell shrinkage and pH_i_ decreases that were blocked by TMEM16A inhibitors NFA, T16A_inh_-A01, CaCC_inh_-A01 (**Figure 2G**). CCh-induced responses were intact in cells from CF patients (**Figure 2H**). Activation of TMEM16A with a pharmacological activator (E_act_; 25 μM) was sufficient to restore both Cl^−^ (shrinkage) and HCO_3_^−^ (pH_i_) secretion responses to VIP in cells from CF patients (**Figure 2H**). In summary, our data suggest serous cell shrinkage during VIP stimulation reflects secretion of both Cl- and HCO3-directly through CFTR (**Figure 2I**).

### NPY reduces CFTR-mediated serous cell fluid and HCO_3_^−^ secretion during VIP stimulation

Beyond the histological observations described above regarding NPY in airways, we also noted that Calu-3 cells, a bronchial adenocarcinoma line frequently used as a serous cell surrogate due to high CFTR and lysozyme expression, express relatively high amounts of NPY1R relative to other airway cancer lines according to public gene expression databases (**Supplemental Tables 1 and 2**). This may be an artifact of Calu-3 cells being cancer cells, but we decided to test for NPY receptor function in primary serous cells.

We observed no secretory responses to 100 nM NPY (**Figure 3A**), but the magnitude of VIP-evoked pH_i_ decreases and cell shrinkage were reduced after NPY (**Figure 3A-B**). As a control, a scrambled NPY peptide had no effect (**Figure 3B**). We hypothesized that G_i_-coupled NPY receptors might blunt the magnitude of VIP-evoked cAMP increases, thus reducing Cl^−^ and HCO_3_^−^ efflux through CFTR. We measured cellular Cl^−^ permeability using 6-methoxy-N-(3-sulfopropyl)quinolinium (SPQ), a dye quenched by Cl^−^ but not by NO_3_^−^ (48, 92). Substitution of extracellular Cl^−^ for NO_3_^−^ results in electroneutral influx of NO_3_^−^ and efflux of Cl^−^, causing a decrease in intracellular [Cl^−^] ([Cl^−^]_i_) and increase in intracellular SPQ fluorescence. Because most Cl^−^ channels are nearly equally permeable to Cl^−^ and NO_3_^−^, the rate of fluorescence increase is roughly equivalent to the anion permeability (93). In the presence of VIP, SPQ fluorescence rapidly increased upon NO_3_^−^ substitution. This was reduced by half in cells stimulated in the presence of 100 nM NPY but not 100 nM scrambled NPY (**Figure 3C-D**). In the presence of CFTR_inh_172, anion permeability was markedly reduced and NPY had no effects (**Figure 3D**), suggesting that NPY directly reduces VIP-stimulated CFTR permeability.

**Figure 3:**
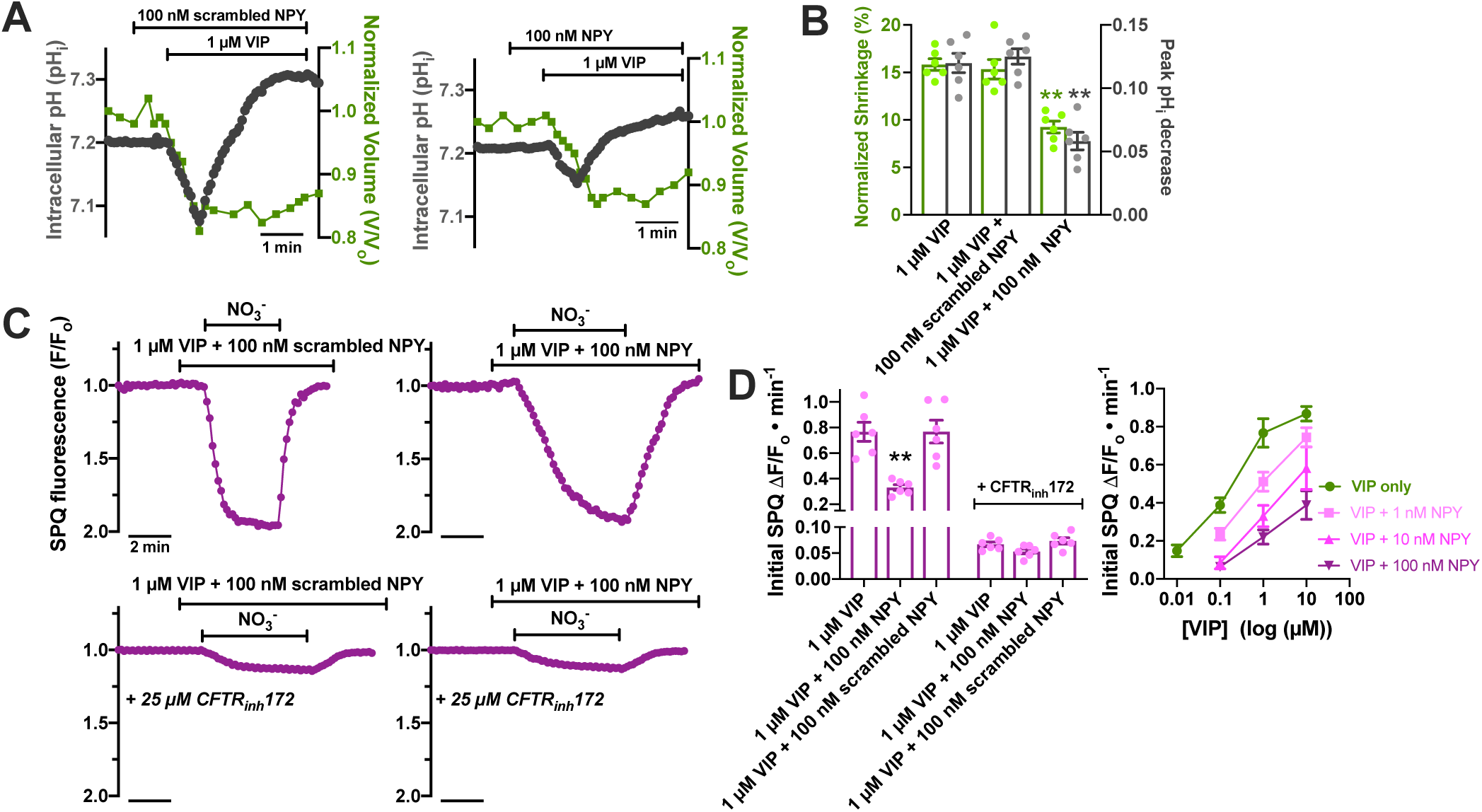
NPY reduces secretory response to VIP and directly reduces anion permeability though CFTR in primary nasal gland serous cells. **(A).** Representative traces showing cells stimulated with VIP in the presence of scrambled NPY (left) or NPY (right). **(B).** Bar graph showing mean ± SEM; cells stimulated with VIP in the presence of NPY exhibited reduced shrinkage (Cl^−^ secretion) and initial acidification (HCO_3_^−^ secretion). Significance determined by 1-way ANOVA with Dunnett’s posttest (VIP only as control group); ** = *p*<0.01 vs control. **(C).** Representative NO_3_^−^ substitution experiments showing changes in SPQ fluorescence with substitution of Cl^−^_o_ for NO_3_^−^_o_, which causes electroneutral exchange of Cl^−^ for NO_3_^−^, a decrease in [Cl^−^]_i_, and a change SPQ fluorescence. The rate of SPQ fluorescence change reflects the relative plasma membrane anion permeability. A downward deflection equals a decrease in [Cl^−^]_i_. (D). Bar graph (left) showing initial rate of SPQ fluorescence (mean ± SEM) change after VIP stimulation, which was inhibited by NPY but not scrambled NPY. In the presence of CFTR_inh_172 (10 μM), rates of SPQ fluorescence change were reduced ∼10-fold and there was no effect of NPY. Right shows rates of SPQ fluorescence change over a range of VIP and NPY concentrations, showing dose dependency of VIP activation of anion permeability and NPY inhibition of anion permeability. *A* and *C* show representative traces, while *B* and *D* show data from at least 6 experiments using acinar cells from 3 patients (2 experiments per patient).

CFTR is activated by PKA downstream of cAMP. We imaged cAMP changes in nasal serous cells in real time using a fluorescent mNeonGreen-based cAMP biosensor (cADDis (94)). 1 μM VIP induced a rapid and reversible increase in cAMP (decrease in cADDis fluorescence) that was blocked by VIPR antagonist VIP_6-28_ (1 μM; **Figure 4A-B**). The cAMP increase was independent of Ca^2+^, as it was not blocked by intracellular and extracellular calcium chelation (**Figure 4C**). Interestingly, we also found no differences in the ability of VIP to increase cAMP in Wt or CF cells (**Supplemental Figure 3**), in contrast to previous hypotheses that cAMP signaling may be defective in CF cells (95). However, NPY (100 nM) significantly reduced the cAMP responses to 0.5 μM and 5 μM VIP (**Figure 4D-E**); the effects of NPY were eliminated in the presence of a NPY1R antagonist BIBO 3304 (5 μM) or in cells treated with pertussis toxin (PTX), which ADP-ribosylates and inactivates G_i_ proteins (**Figure 4D-E**). These data demonstrate that NPY reduces cellular anion efflux through CFTR to blunt Cl^−^, HCO_3_^−^, and fluid secretion from these cells.

**Figure 4:**
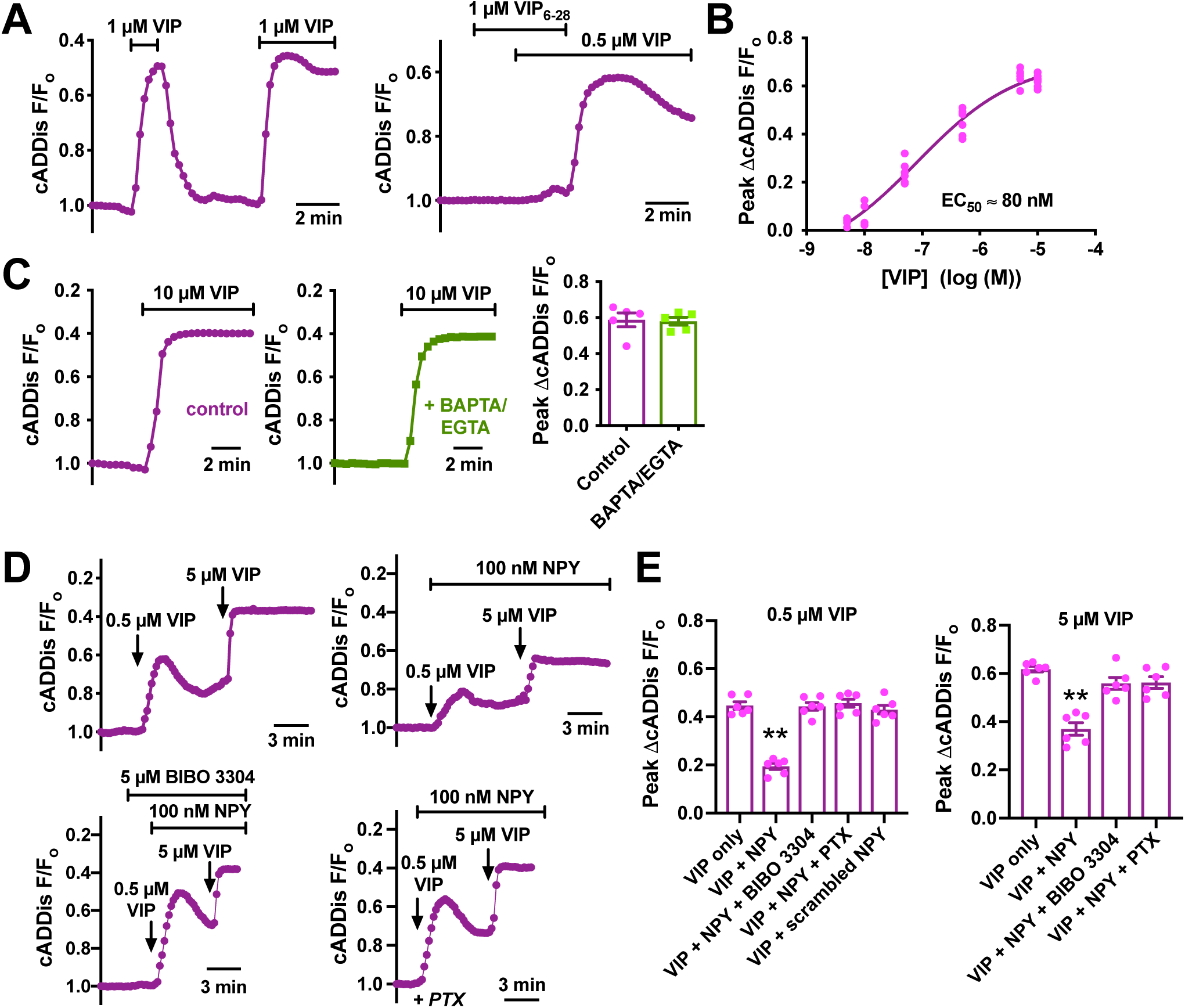
NPY inhibits VIP-induced cAMP increases in primary nasal gland serous cells. **(A).** Representative traces of cADDis fluorescence (upward deflection of trace = increase in cAMP) showing reversible VIP-activated cAMP increases blocked by VIP receptor antagonist VIP_6-28_. **(B).** Dose response showing peak cADDis fluorescence changes with VIP. Each data point is a separate experiment; graph shows data from at least 3 serous cells from at least 3 patients (at least one per patient) for each concentration. **(C).** Representative traces (left) and bar graph (right) showing lack of inhibiton of cADDis responses by calcium chelation (10 μM BAPTA-AM loading for 30 and stimulation in solution containing no added calcium + 1 mM EGTA). Bar graph shows mean ± SEM of 5 experiments from serous cells from 2 different patients. No significant difference by Student’s *t* test. **(D).** Peak cAMP responses to 0.5 μM and 5 μM VIP (top left) were inhibited by NPY (top right); NPY reduction of cAMP responses were abolished by NPY1R antagonist BIBO 3304 (bottom left) or pretreatment with pertussis toxin (PTX). **(E).** Bar graphs showing mean ± SEM of peak responses from experiments as in *D* at 2 different VIP concentrations. Shown are data points from at least 6 experiments using serous cells from at least 3 patients (at least 2 experiments per patient). Significance determined by 1-way ANOVA with Dunnett’s post test (VIP only as control); ** = *p*<0.01 vs VIP only.

To facilitate polarized studies of serous cells, we used previously published culture methods for gland acinar cells that preserve a serous phenotype (96-98). Serous cells cultured at air liquid interface (ALI) expressed serous marker Muc7 (99), VIP1R, and VIP2R by Western (**Figure 5A**). Mucous maker Muc5B was not detected (**Figure 5A**). Serous cell markers lysozyme (100, 101), Muc7, and VIPR1 and VIPR2 were detected by immunofluorescence (**Figure 5B-C**). Many of these same markers were detected in Calu-3 cells (**Supplemental Figure 4-5**). We used ELISAs to confirm that serous cell cultures expressed serous cell Muc7 but not goblet cell mucin Muc5AC or mucous cell mucin Muc5B (**Supplemental Figure 6**).

**Figure 5:**
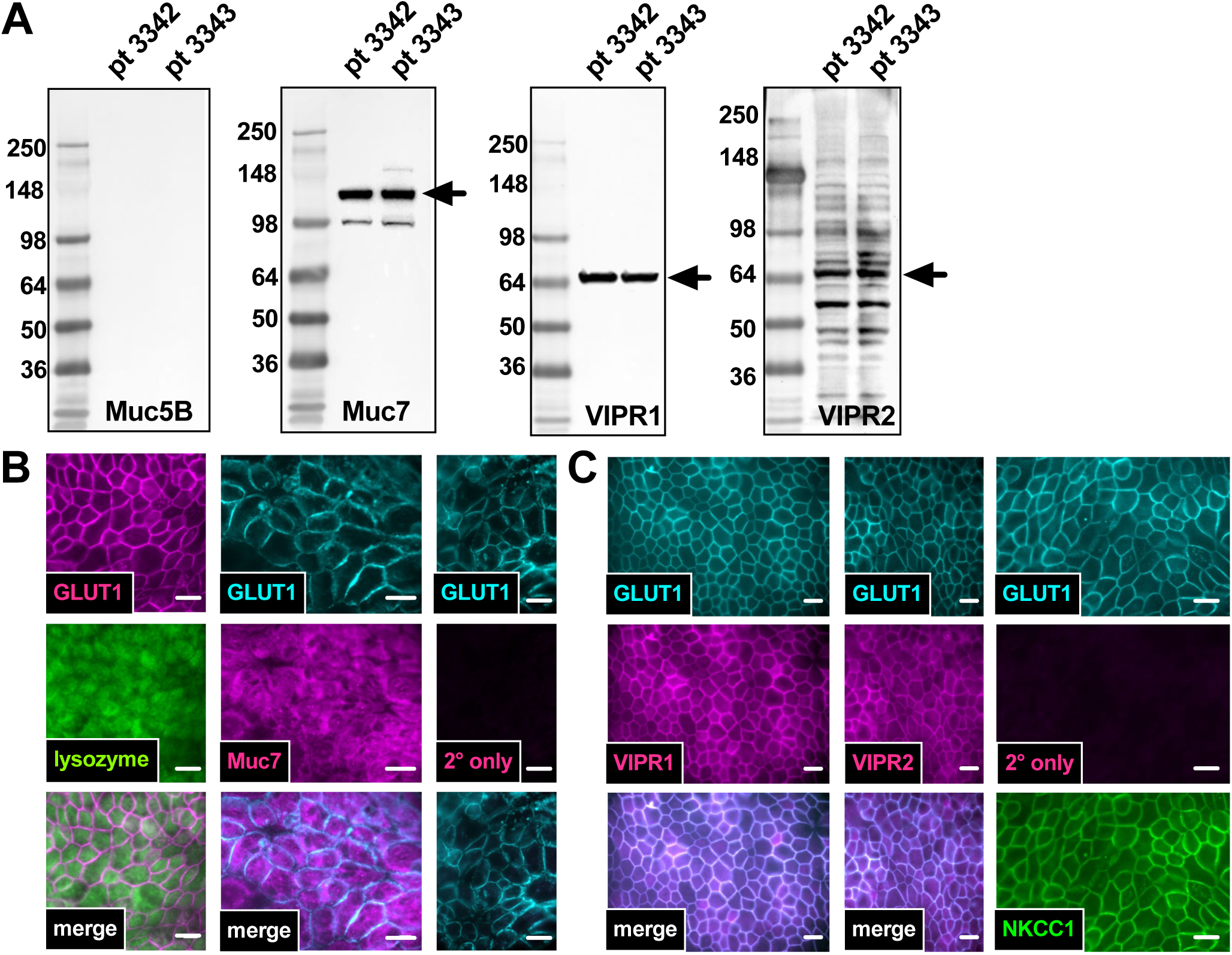
Expression of serous cell markers by primary nasal serous cell ALI cultures. **(A).** Acinar cells isolated from middle or inferior turbinate were cultured as indicated in the text. ALIs were subject to Western Blot for mucous cell marker Muc5B, serous cell marker Muc7, and VIP receptors VIPR1 and VIPR2. Results from cultures from two different patients are shown, representative of results observed from cultures at least 3 independent experiments. **(B).** Fixed cultures were immune-stained for serous cell markers lysozyme and Muc7, which showed punctate cytoplasmic staining similar to serous-like secretory granules. **(C).** Immunocytochemistry for VIPR and VIPR2 revealed basolateral staining similar to that observed with GLUT1 and NKCC1. All images in *B* and *C* are representative of results observed in cultures from at least 3 separate patients. Scale bars are 20 μm.

**Figure 6:**
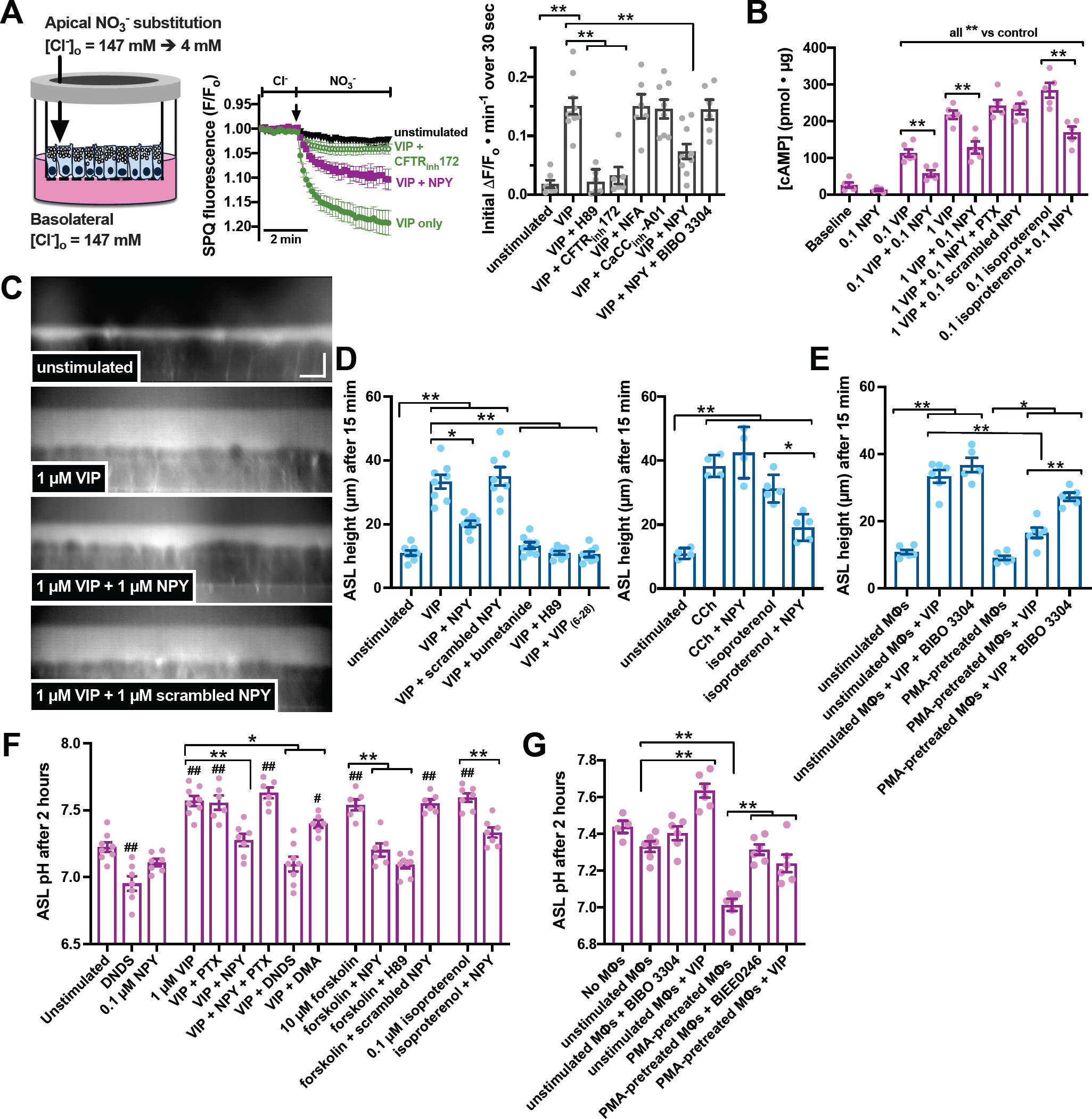
Modulation of fluid and HCO_3_^−^ secretion by VIP and NPY in serous cell ALI cultures. **(A).** Apical NO_3_^−^ substitution experiments (representative traces, left) and rates of SPQ change (bar graph right) during stimulation with VIP ± NPY in the presence or absence of indicated inhibitors. **(B).** ELISA results from steady-state cAMP measurements during stimulation with VIP or isoproterenol ± NPY or scrambled NPY. Concentrations shown are μM. Pertussis toxin (PTX) was used to demonstrate NPY effects were dependent on G_i_ signaling. **(C).** Representative orthogonal views of Texas red dextran-labeled ASL in primary serous cell ALIs; scale bar is 10 μm in both *x* and *z* direction. **(D).** ASL height after 15 min basolateral stimulation as indicated. **(E).** ASL height in ALIs incubated in the presence of MΦs and MΦ-conditioned media with basolateral compounds as indicated. **(F).** ASL pH measured using SNARF-1-dextran in cultures stimulated as indicated for 2 hours. Concentrations shown are μM. **(G).** ASL pH in ALIs incubated in the presence of MΦs and MΦ-conditioned media with basolateral compounds as indicated. All bar graphs show mean ± SEM of at 6 independent experiments using ALI cultures from at least 3 patients (2 cultures per patient). Significance in each bar graph determined by 1-way ANOVA with Bonferroni posttest; *= *p* <0.05 and ** = *p* <0.01 vs bracketed groups and ## and # in *F* represent *p* <0.05 and *p* <0.01, respectively vs unstimulated conditions.

Serous cell ALIs also expressed functional apical CFTR; when ALIs were loaded with SPQ, apical substitution of Cl^−^ for NO_3_^−^ led to a decrease in [Cl^−^]_i_ (increase in SPQ fluorescence) that was enhanced by VIP (1 μM), blocked by CFTR_inh_172 (10 μM), and blunted in the presence of NPY (100 nM; **Figure 6A**). Similar to studies of freshly isolated cells above, TMEM16A inhibitors did not affect VIP-activated Cl**^−^** permeability (**Figure 6A**). ALIs were resistant to viral expression of cADDis, but we measured changes in steady-state cAMP levels 5 min after stimulation with VIP ± NPY. NPY reduced cAMP increases in response to VIP or isoproterenol, and this was abrogated by PTX (**Figure 6B**), suggesting effects of NPY on cAMP are dependent on activation of G_i_-coupled receptors.

Airway surface liquid (ASL) was labeled with Texas red dextran and imaged with confocal microscopy to track fluid secretion in serous cell ALIs stimulated VIP (1 μM) ± NPY (100 nM) on the basolateral side. VIP increased ASL height, and this was inhibited by NKCC1 inhibitor bumetanide (100 μM), PKA inhibitor H89 (10 μM), or VIPR antagonist VIP_(6-28)_ (1 μM) (**Figure 6C-D**). NPY, but not scrambled NPY, inhibited VIP-induced secretion (**Figure 6C-D**). Ca^2+^-driven 100 μM CCh-induced secretion was unaffected by NPY (**Figure 6D**), while effects of another cAMP-elevating agonist, isoproterenol, was inhibited by NPY (**Figure 6D**), showing effects of NPY were specific for cAMP-elevating agonists.

Primary human monocyte-derived macrophages (MΦs) stimulated with PKC-activating phorbol myristate acetate (PMA; 100 nM for 48 hrs) produce NPY ((62) and **Supplemental Figure 7**), were washed to remove PMA and incubated for 24 hours in a 24 well plate. Serous cells on transwells were then transferred into the same plates above the MΦs in the 24 hour conditioned media on the basolateral side. Addition of VIP (2 μM) caused an increase in ASL height that was reduced in the presence of PMA-stimulated MΦs compared with unstimulated MΦs (**Figure 6E**). In the presence of PMA-stimulated MΦs, VIP-induced fluid secretion was increased by addition of NPY1R antagonist BIBO 3304 (1 μM).

**Figure 7:**
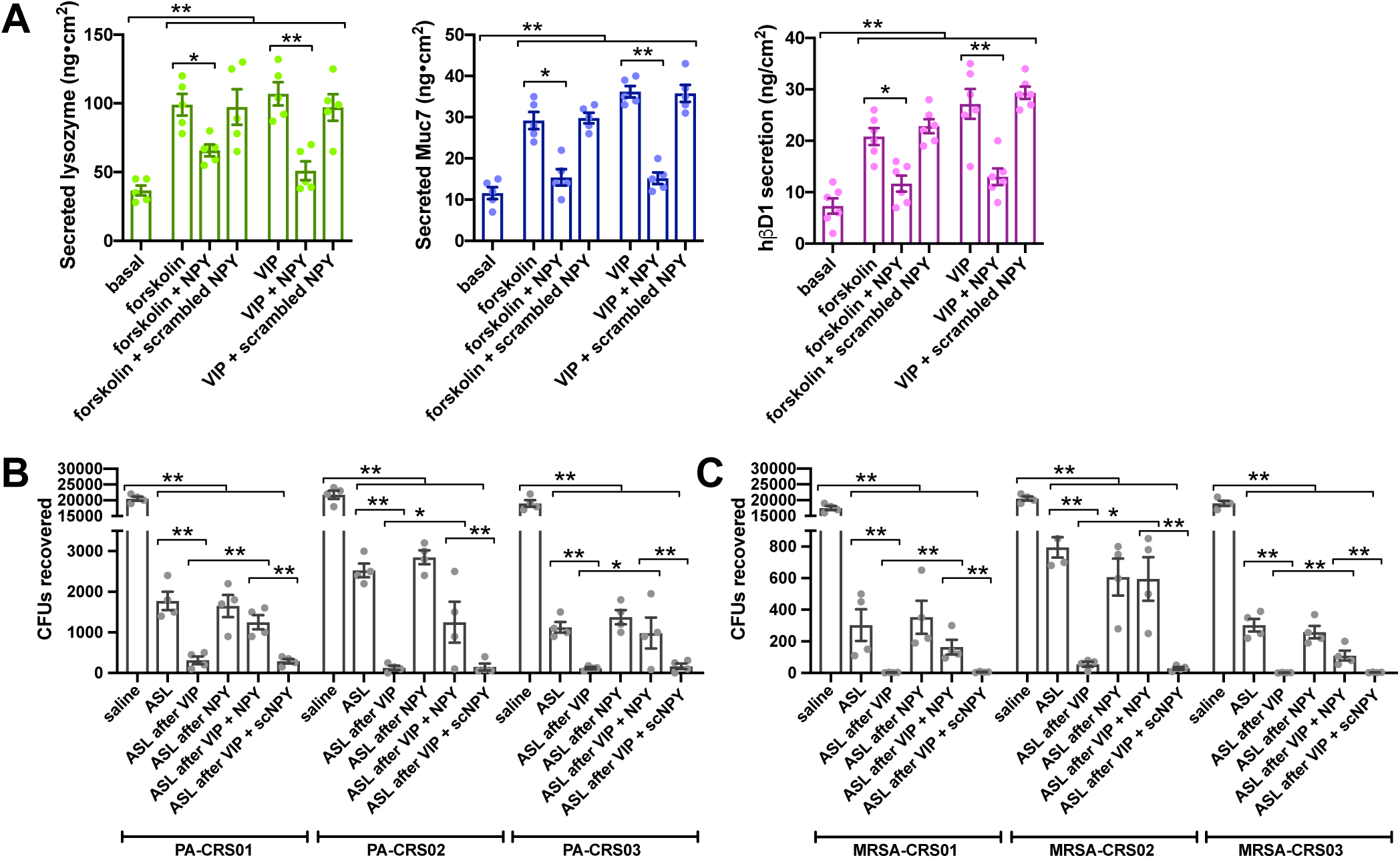
Antimicrobial peptide secretion and antibacterial efficacy of serous cells secretions are enhanced by VIP but reduced by NPY. **(A).** ALIs were stimulated basolaterally for 2 hours in the presence of forskolin (10 μM) or VIP (1 μM) ± NPY (100 nM) or scrambled NPY (100 nM) as indicated. ASL was collected by washing the apical surface with 25% saline and assayed for lysozyme, Muc7, and hβD1 by ELISA. Results shown are mean ± SEM from at least 3 ALIs from at least 3 individual patients. **(B-C).** ASL from similar experiments was mixed with strains of *P. aeruginosa* (*B*) or methicillin-resistant *S. aureus* (MRSA; *C*) isolated from CRS patients followed by incubation (37°C; 5% CO_2_) and plating for CFU counting as indicated in the text. Bar graphs show mean ± SEM of at least 5 experiments using ALIs from at least 3 different patients; ** and * indicate *p*<0.01 and *p*<0.05, respectively, between bracketed groups. Significance determined by 1-way ANOVA with Bonferroni posttest.

ASL was labeled with SNARF-1-dextran, a pH probe with ratiometric emission (580 and 650 nm) and thus insensitive to changes in volume. SNARF-1-dextran was sonicated in perfluorocarbon, allowing measurement of pH within the physiological ASL with no addition of extra fluid (102). Steady-state unstimulated ASL pH was 7.2 ± 0.04, equivalent to a [HCO_3_^−^] of 15 mM by Henderson-Hasselbach with 5% CO_2_ ([HCO_3_^−^]_i_ = 1.2 mM × 10^pH-6.1^). ASL pH was reduced to 6.9 ± 0.06 (equivalent to 7.6 mM HCO_3_^−^) by NBC inhibitor 4,4’-dinitrostilbene-2,2’-disulfonic acid (DNDS; 30 μM) but was not significantly reduced with NPY (100 nM) (**Figure 6F**). VIP (1 μM) increased ASL pH to 7.6 ± 0.04 (equivalent to 38 mM HCO_3_^−^), suggesting VIP stimulated HCO_3_^−^ secretion. VIP-increased ASL pH was reduced by NPY (7.3 ± 0.05) or DNDS (7.1 ± 0.03). Effects of NPY were blocked by PTX. NPY had similar inhibitory effects on ASL pH increases with forskolin and isoproterenol (**Figure 6F**). Note that with increase in ASL volume (**Figure 6C**) as well as buffering capacity, the amount of secreted HCO_3_^−^ would be even greater than predicted changes in [HCO_3_^−^].

Serous cell ALIs were incubated in the presence of unstimulated or PMA-stimulated MΦs as above and ASL pH was measured 2 hours later. ASL pH was not different in the presence or absence of unstimulated MΦs, but PMA-stimulated MΦs reduced steady-state ASL pH (**Figure 6G**). This effect was inhibited by an NPY1R antagonist (BIBO 3304; 1 μM) and pH was also increased by addition of VIP (1 μM) (**Figure 6G**). Effects of NPY on HCO_3_^−^ secretion were verified using a real-time HCO_3_^−^ secretion assay using larger apical volumes of SNARF-1-dextran, which confirmed secretion was dependent on apical CFTR (**Supplemental Figure 8**).

**Figure 8:**
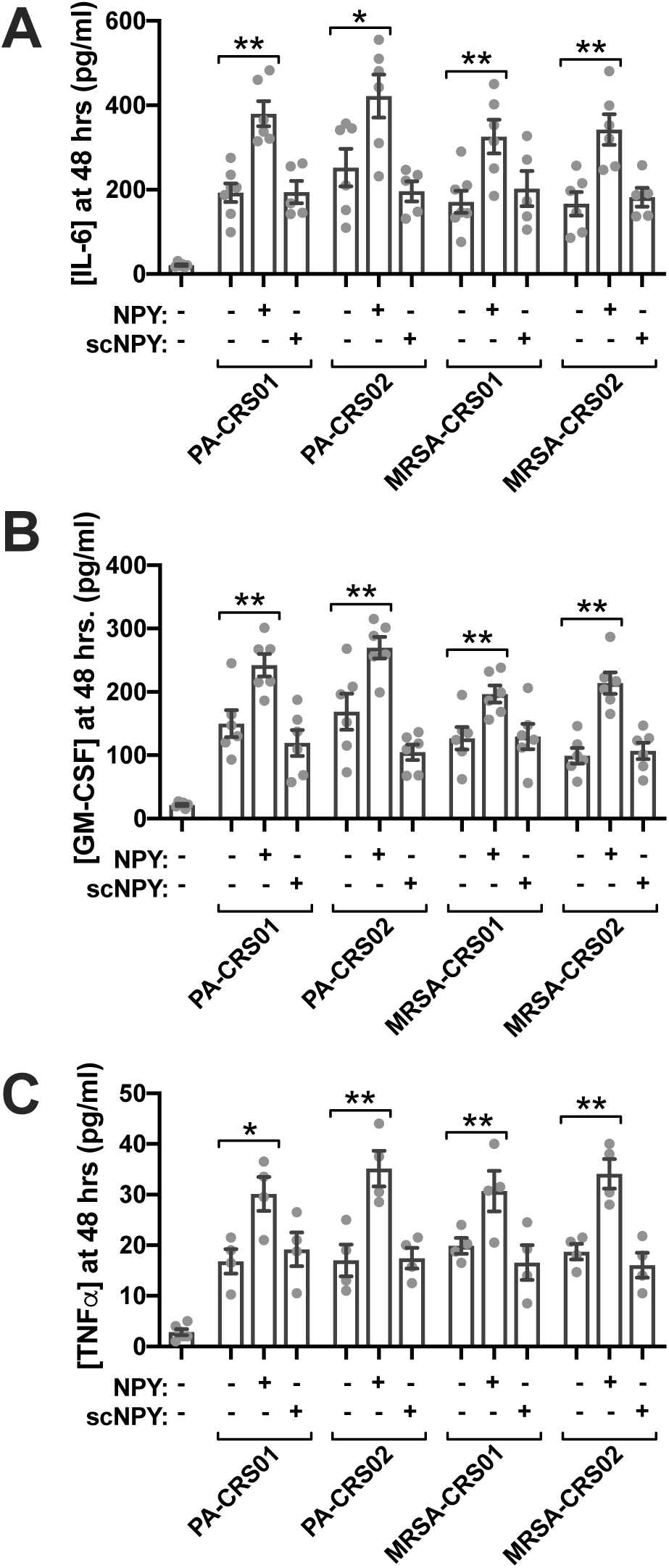
Serous cell cytokine secretion in response to clinical bacteria strains is increased by NPY. Primary serous cell ALIs were treated apically with heat-killed bacteria, followed by 48 hr incubation ± basolateral NPY (100 nM) or scrambled NPY (scNPY; 100 nM). Basolateral media was collected for quantification of IL-6 (*A*), GM-CSF (*B*), and TNFα (*C*). Bar graphs shown mean ± SEM of at least 5 experiments using cells grown from at least 3 different patients. Significance determined by 1-way ANOVA with Bonferroni posttest (comparing the three bars for each separate strain); ** p<0.01 and * p<0.05 between bracketed bars.

### NPY inhibits VIP-evoked increases in serous cell antimicrobial peptide secretion

Serous cells secrete a variety of antimicrobial peptides, and secretion can involve cAMP. It is likely that the same stimuli that activate fluid secretion likely activate protein secretion, which is also driven by Ca^2+^ and cAMP (103, 104). To test if NPY can reduce VIP-induced secretion of antimicrobials, we measured secreted levels of lysozyme, Muc7, and β-defensin 1 (hβD1). Cells were stimulated basolaterally with forskolin (5 μM) or VIP (1 μM) in the presence or absence of NPY or scrambled NPY (100 nM). Forskolin and VIP both increased secretion of lysozyme, Muc7, and βD1, and this was reduced by NPY (**Figure 7A**).

### VIP increases bactericidal activity of serous cell secretions while NPY reduces it

Carbonate and/or HCO_3_^−^ has been reported to enhance antimicrobial activity of airway antimicrobial secretions (45, 105). We did observe a small effect of HCO_3_^−^ on antimicrobial activity of secretions produced by Calu-3 bronchial serous-like cells (**Supplemental Figure 9**). However, we hypothesized that NPY might have more profound effects on antimicrobial activity through inhibition of both HCO_3_^−^ secretion and antimicrobial peptide secretion. We tested the anti-bacterial efficacy of apical washings of serous cells stimulated with VIP ± NPY or scrambled NPY. VIP increased the antibacterial effects of ASL washings (as measured by CFU counting) against both clinical isolates of gram negative *P. aeruginosa* (**Figure 7B**) and methicillin resistant gram positive *S. aureus* (MRSA; **Figure 7C**), fitting with data above suggesting increased antimicrobial peptide secretion. Addition of NPY itself had no effect, but NPY substantially blunted the effect of VIP against either species of bacteria (**Figure 7B-C**). Scrambed NPY did not reduce the increased efficacy observed with VIP (**Figure 7B-C**). A fluorescent live-dead assay (Syto9 and propidium iodide staining) confirmed reduced efficacy of NPY+VIP stimulated ASL after only 5 min incubation with *P. aeruginosa* (**Supplemental Figure 10**).

**Figure 9:**
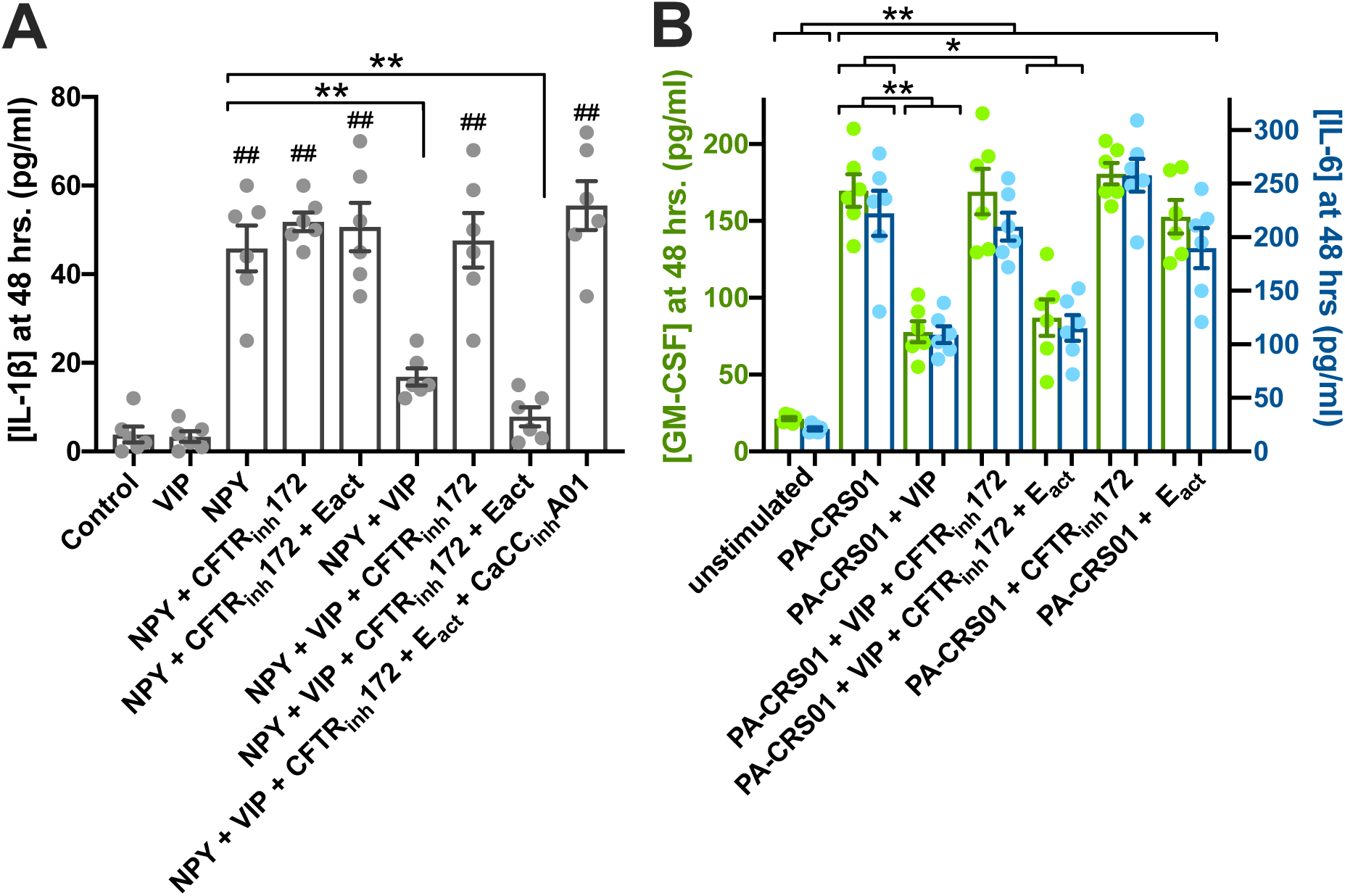
Anti-inflammatory effects of VIP require CFTR conductance, but can be restored by TMEM16A activation. Primary serous cell ALIs were treated basolaterally with VIP (100 μM) and/or NPY (100 nM) and treated apically with CFTR inhibitor CFTR_inh_172 (15 μM), TMEM16A activator E_act_ (15 μM), and/or TMEM16A inhibitor CaCC_inh_-A01 (15 μM). Basolateral media was collected after 48 hrs. and assayed for IL-1β. **(B)** Primary serous cell ALIs were treated apically with heat-killed *P. aeruginosa*, followed by 48 hrs incubation ± basolateral NPY and/or VIP as well as ± apical CFTR_inh_172 and/or E_act_. Basolateral media was collected after 48 hrs. and assayed for GM-CSF or IL-6.

### NPY has pro-inflammatory effects in primary serous acinar cells

Both VIP and NPY have immunomodulatory roles in many tissues (106-109), including VIP having anti-inflammatory or protective effects in parotid acini (106, 110-113) and NPY having pro-inflammatory effects in leukocytes (85). Acinar cells from parotid and pancreatic exocrine glands can make and release cytokines (114-117). Infection of isolated human tracheal submucosal gland cells with rhinovirus, which can activate TLR3 (118), increases IL-1α, IL-1β, IL-6, and IL-8 (119). TLR4 is also expressed in pig tracheal acinar cells (120), and submucosal TLR4 levels may be elevated in CF (121). We hypothesized that airway gland cells may be an overlooked significant contributor to the airway cytokine milieu, and this may be modulated by VIP and/or NPY.

In primary nasal serous cell ALIs, the TLR4 activator lipopolysaccharide (LPS) and TLR3 activator poly(I:C) induced secretion of IL-6, TNFα, IL-1β, and granulocyte macrophage colony stimulating factor (GMCSF (**Supplemental Figure 11**). TLR2 activator lipotechoic acid (LTA) also increased secretion of IL-6 and TNFα, while TNFα itself increased secretion of IL-1β and GM-CSF (**Supplemental Figure 11**). Furthermore, a type 2 inflammatory cocktail of IL-4 and IL-13 (122) also increased secretion of GM-CSF (**Supplemental Figure 11**). Airway gland serous cells can thus respond to and secrete a variety of inflammatory cytokines.

While NPY or VIP had no effect alone on IL-6, TNFα, or GMCSF, NPY increased IL-1β production ~2-fold and significantly increased cytokine production (~50%) during LPS, LTA, IL-4+IL-13, and TNF-α (**Supplemental Figure 11)** stimulation. Effects of NPY were blocked by pertussis toxin, implicating a GPCR G_i_-coupled pathway. In contrast, VIP slightly reduced cytokine secretion (25-50%) when combined with inflammatory stimuli, while these reductions were eliminated in the presence of NPY (**Supplemental Figure 11**). Together, these data suggest that VIP has a small anti-inflammatory effect while NPY is pro-inflammatory when combined with a broad range of stimuli.

A strong Th2 environment by itself may increase other inflammatory responses in airway cells (123). Co-stimulation with IL-4+IL-13 increased IL-6 and GM-CSF secretion in response to either poly(I:C) or LPS, and this was enhanced further in the presence of NPY (**Supplemental Figure 12A**), suggesting that NPY is pro-inflammatory even within the context of elevated IL-4 and IL-13 in inflammatory airway diseases. To validate results from cultured cells, we incubated freshly dissociated primary serous celsl seeded at high density with TNFα or poly(I:C) ± NPY or scrambled NPY for 18 hours. TNFα and poly(I:C) increased secretion IL-33, GM-CSF, or IL-6, and this was enhanced by NPY but not scrambled NPY (**Supplemental Figure 12B**), supporting that airway gland serous cells can secrete several cytokines involved in allergy, asthma, and chronic rhinosinusitis, and confirming NPY is pro-inflammatory.

We examined cytokine release in response to heat-killed clinical CRS isolates of gram negative *P. aeruginosa* and gram positive methicillin-resistant *Staphylococcus aureus* (MRSA). Incubation of serous cell ALIs with either species of bacteria increased secretion of IL-6, GM-CSF, and TNFα (**Figure 8**). NPY (100 nM), but not scrambled NPY, significantly increased cytokine release, supporting that NPY has pro-inflammatory effects during airway infections.

### Anti-inflammatory effects of VIP require apical functional CFTR conductance, but activation of TMEM16A can substitute for CFTR

In airway cells, Cl^−^ conductance has been suggested to be anti-inflammatory (124, 125), with increased intracellular [Cl^−^]_i_ promoting inflammation (126). In serous cells stimulated with VIP, [Cl^−^]_i_ may be higher if there is a lack of apical CFTR efflux pathway in CF patients. To test if CFTR was required for anti-inflammatory effects observed with VIP, we first stimulated serous cell ALIs with NPY, which increased IL-1β secretion; NPY-induced IL-1β was not altered by CFTR_inh_172 or activation of TMEM16A (E_act_) (**Figure 9A**). However, VIP reduced IL-1β secretion by >50% (**Figure 9A**). CFTR_inh_172 reversed the anti-inflammatory effect of VIP, while adding E_act_ restored the anti-inflammatory effect of VIP (**Figure 9A**). The effect of E_act_ reversed with CaCC_inh_-A01 (**Figure 9A**). These data suggest that CFTR is required for the anti-inflammatory effects of VIP but TMEM16A can substitute. However, the Cl^−^ conductance itself is not sufficient, as E_act_ did not have anti-inflammatory effects in the absence of VIP. This may be because a reduction in [Cl^−^]_i_ would require counter-ion (K^+^) flux that would be activated downstream of a cAMP-secretagogue like VIP, as we have previously suggested through cAMP-activated Ca^2+^ signals (11), but not during direct activation of TMEM16A with apical E_act_. We saw similar results when serous cells were stimulated with heat-killed *P. aeruginosa*. VIP reduced GM-CSF and IL-6 secretion, but these effects were blocked by CFTR_inh_172 and subsequently restored by E_act_ (**Figure 9B**). CFTR_inh_172 and E_act_ had no effect alone on *P. aeruginosa*-induced GM-CSF or IL-6 secretion (**Figure 9B**), again suggesting that an apical Cl^−^ conductance is necessary, but not sufficient, for anti-inflammatory effects of VIP.

## DISCUSSION

This paper reveals several important insights into airway gland serous cell physiology. First, we directly demonstrate that serous cells secrete HCO_3_^−^ in addition to Cl^−^ during VIPergic stimulation. Our experiments suggest this is conductive HCO_3_^−^ efflux through CFTR with little contribution from Cl^−^/HCO_3_^−^ exchangers such as pendrin. We found no defect in cAMP signaling in primary serous cells, supporting that appropriate pharmacological correction of mutant CFTR function (127) would restore fluid secretion in response to appropriate physiological stimuli (e.g., VIP). However, in patients that cannot benefit from CFTR correction (e.g., those with a premature stop code-on mutation), our data suggest activation of TMEM16A, bypassing CFTR, is sufficient to restore HCO_3_^−^ efflux during VIP stimulation in CF serous cells (128).

We also demonstrate a novel inverse relationship between NPY and VIP in the regulation of serous cell secretion. Our data here suggest that VIP may promote watery secretions of glands through elevated fluid and HCO_3_^−^ secretion to thin mucus. However, we hypothesize that under conditions of increased NPY (e.g., in asthma), the ability of VIP to stimulate fluid and HCO_3_^−^ secretion is markedly impaired, as is the secretion of antimicrobial peptides. Coupled with increased inflammation in the presence of NPY, our data suggest NPY may have multiple detrimental effects in diseases of mucus thickening/mucostasis. We (129) and others (130-132) have shown that NPY decreases airway ciliary beat frequency, which may further impair mucociliary clearance.

Patients challenged with allergens produce nasal secretions that have detectible levels of VIP (31, 32), suggesting this peptide is released in large amounts during the airway allergic response, possibly through histamine activation of sensory neurons (37). Allergic rhinitis patients may have a higher density of sinonasal VIPergic fibers (21, 22, 33-35), increased VIP receptor expression (36), and baseline nasal secretions with elevated concentrations of VIP compared with control individuals (32). This may thin out mucus by increasing secretion of HCO_3_^−^ and fluid from gland serous cells. In contrast, elevations of NPY may thicken mucus in some asthma patients, with the balance of these two peptides contributing toward setting airway mucus rheology. We hypothesize that in some asthma, COPD, or CRS patients, NPYR1 antagonists may be useful that to thin secreted mucus, enhance antimicrobial secretion, and reduce inflammatory responses from gland acini by relieving repression of VIP-induced signaling.

The important contribution of exocrine acinar cells to inflammation is already established in parotid and pancreatic acini within the context of Sjögren’s syndrome and pancreatitis, respectively (114-116). However, this has been largely unstudied in the airway. In bronchi, gland volume may be up to 50-fold larger than the volume of surface goblet cells (5, 6, 133-135). Gland acini are likely significant contributors to the airway cytokine milieu, particularly when barrier dysfunction occurs during chronic inflammation in CRS, COPD, asthma, or CF (136-138) and/or when gland hypertrophy and hyperplasia occur during COPD and asthma (25, 27, 28, 30). Elevations of NPY may alter submucosal gland function by both reducing cAMP-driven CFTR-mediated secretion as well as enhancing production of cytokines like GM-CSF and IL-1β that are important in allergic inflammation (139-142), airway neutrophil or eosinophil infiltration (143, 144), and Th2 polarization (145-147). NPY by itself increase IL-1β secretion, and IL-1β polymorphisms may contribute to CF (148) or CRS (149); it remains to be determined if these polymorphisms relate to expression or secretion of IL-1β from gland acini. Regardless, NPY-increased serous cell-derived cytokines likely help to drive inflammation.

Finally, our data support previous observations (124, 125) that the Cl^−^ channel activity of CFTR is anti-inflammatory during VIP stimulation. A loss of these anti-inflammatory effects of VIP in CF patients lacking functional CFTR may contribute to the hyperinflammatory phenotypes reported (150). As we saw for Cl^−^ and HCO_3_^−^ secretion, our data suggest that activation of TMEM16A can also compensate for loss of CFTR to restore anti-inflammatory effects of VIP, suggesting another possible benefit to targeting TMEM16A in CF submucosal glands of patients who cannot benefit from CFTR potentiator and/or corrector therapies due to CFTR genotype.

## METHODS

### Experimental Procedures

Isolation of primary serous acinar cells, immunofluorescence, and live cell imaging of acinar cell volume, pH_i_ (SNARF-5F), and Cl^−^ (SPQ) was carried out as described (10-12, 47, 48). ASL height and pH measurements and ELISAs were carried out as previously reported (92, 129, 151-157). Bacterial growth assays were carried out as previously described (158, 159). More detailed methods for all procedures as well as specific reagents used are provided in the **Supplemental Materials**.

### Study Approval

Tissue was acquired with IRB approval (University of Pennsylvania protocol # 800614) in accordance with the University of Pennsylvania School of Medicine guidelines regarding residual clinical material in research, the United States Department of Health and Human Services code of federal regulations Title 45 CFR 46.116, and the Declaration of Helsinki.

### Serous cell isolation and culture

Primary human nasal serous acinar cells were used to study Cl^−^/fluid and HCO_3_^−^ secretion. Studies of human turbinate submucosal gland serous cells are directly relevant to the understanding of mechanisms of CRS, particularly CF-related CRS (160), and turbinate gland serous cells approximate gland serous cells from the lower airway. Histology suggests that nasal airway glands are similar to tracheal/bronchial glands (161), and we have established that pig bronchial serous cell responses are identical to human turbinate serous cells (11, 12). Working with human cells has important advantages over mice, as data from intact glands (19, 162, 163) and our own studies (10-12, 46-48) have established important differences between mouse serous cells and those from pigs and humans.

Patients undergoing medically indicated sinonasal surgery were recruited from the Department of Otorhinolaryngology at the University of Pennsylvania with written informed consent as previously described (151-154). Inclusion criteria were patients ≥18 years of age undergoing surgery for sinonasal disease (CRS) or other procedures (e.g., trans-nasal approaches to the skull base) where tissue was classified as “control.” Exclusion criteria included history of systemic inheritable disease (e.g., granulomatosis with polyangiitis or systemic immunodeficiencies) with the exception of cystic fibrosis (CF). Members of vulnerable populations were not included.

Comparisons made here between non-CF and CF cell Cl^−^ and HCO_3_^−^ secretion are valid because SNARF and SPQ properties were identical between CF and non-CF cells, and both genotypes had identical resting [Cl^−^]_i_, resting pH_i_, and intracellular pH_i_ buffering capacity (**Supplemental Figures 13-14**).

For culturing, acinar cells were washed with and resuspended in 1:1 MEME plus 20% FBS, 1x pen/strep, gentamycin (100 μg/ml), and amphotericin B (2.5 μg/ml) as described by Finkbeiner (96). Cells were seeded (∼3×10^5^ cells per cm^2^) on transparent Falcon filters (#353095; 0.3 cm^2^; 0.4 μm pores) coated with human placental collagen. After confluence, the media was changed to MEME + Lonza bronchial epithelial cell culture supplements (5 μg/ml insulin, 5 μg/ml transferrin, 0.5 μg/ml hydrocortisone, 20 ng/ml triiodothyronine, 20 nM retinoic acid, 2 mg/ml BSA) but not EGF, with 2% NuSerum. Media lacking EGF combined with the plastic type of these transwell filters was previously shown to differentiate cells into a serous phenotype (96, 164). After 5 days of confluence, TEER reached ∼300 – 500 Ω•cm^2^ and cells were fed with the media above lacking NuSerum on the basolateral side while the apical side was washed with PBS and exposed to air. Cells were used after 2-4 weeks at air-liquid interface.

### Imaging of intracellular cAMP dynamics in isolated nasal gland serous cells

Isolated acinar cells were plated for 30 min on glass coverslips, followed by washing and addition of serum-free Ham’s F12K (Gibco) containing cADDis expressing BacMam (Montana Molecular) plus 5 mM NaButyrate to enhance expression. Cells were imaged after 24 hrs incubation at 37 °C. A BacMam vector was previously used to express proteins in lacrimal gland acinar cells (165-167). Cells were imaged as above under CO_2_/HCO_3_^−^ conditions using a standard GFP/FITC filter set (Semrock) on a Nikon microscope (20× 0.75 Plan Apo objective) equipped with a QImaging Retiga R1 camera and XCite 110 LED illumination system. Data were acquired with Micromanager (168). Experiments were done under ion substitution conditions (high K^+^) to reduce volume changes as previously described (10-12, 47, 48) to ensure that cADDis fluorescence changes were not artifacts of cell volume change during activation of secretion, confirmed by pilot experiments using mNeonGreen-only BacMam. For experiments with pertussis toxin (PTX), PTX was included with the BacMam virus infection reaction (∼24 hours pretreatment).

### Primary culture of human monocyte-derived macrophages (Mϕs)

Monocytes were isolated from healthy apheresis donors by RosetteSep™ Human Monocyte Enrichment Cocktail (Stem Cell Technologies) by the University of Pennsylvania Human Immunology Core and provided as de-identified untraceable cells. Monocytes were differentiated into macrophages by 10 days of adherence culture in high glucose RPMI media containing 10% human serum. Differentiation to Mϕs was confirmed by functional expression of markers including histamine H1 receptors (169, 170) determined by Ca^2+^ imaging (**Supplemental Figure 15**) with specific antagonists as well as secretion of appropriate cytokines in response to M1 vs M2 polarization stimuli (**Supplemental Figure 7**).

### Statistics

Numerical data was analyzed in Microsoft Excel or GraphPad Prism. Statistical tests were performed in Prism. For multiple comparisons with 1-way ANOVA, Bonerroni posttest was used when preselected pairwise comparisons were performed, Dunnett’s posttest was used when values were compared to a control set. Tukey-Kramer posttest was used when all values in the dataset were compared. A *P* value <0.05 was considered statistically significant. All data are mean ± SEM from independent experiments using cells from at least 4 patients.

## Acknowledgements

We thank N. Cohen and L. Chandler (Philadelphia VA Medical Center) for clinical bacteria isolates and P. aeruginosa strains PAO-1 and PAO-GFP, and J. Riley (University of Pennsylvania Department of Microbiology and Human Immunology Core) for access to primary human monocytes. We thank M. Victoria (University of Pennsylvania Department of Otorhinolaryngology) for excellent technical assistance with differentiation of macrophages and molecular biology and B. Chen (University of Pennsylvania Department of Otorhinolaryngology) with assistance growing initial cultures of primary serous cells. This work was supported by grants from the Cystic Fibrosis Foundation (LEER16G0) and National Institutes of Health (R21AI137484, R01DC016309). The sponsors had no role in study design, data collection, interpretation, writing, or the decision to submit.

## Author Contributions

D.B.M., M.A.K., and R.J.L. performed experiments, analyzed data, and interpreted results. M.A.K., C.C.L.T., P.P., N.D.A., and J.N.P. aided with tissue and primary cell acquisition, consenting of patients, maintenance of clinical databases and records, and intellectually contributed to interpretation of the study. R.J.L. conceived the study and wrote the paper with input and approval from all authors.

