## Supplemental Material for "Inverse regulation of secretion and inflammation in human airway gland serous cells by neuropeptides upregulated in allergy and asthma"

### Supplemental Methods

#### *Reagents and Solutions*

Reagents for fluorescence microscopy (SNARF-5F-AM, SNARF-1 dextran, 6-methoxy-N-(3-sulfopropyl)quinolinium [SPQ], Texas Red dextran [10,000 MW], BAPTA-AM) were from Thermo Fisher Scientific. ELISA development kits were from Peprotech (GM-CSF Cat # 900-K30, IL-6 Cat # 900-K16, TNF$\alpha$ Cat # 900-K25, IL-1$\beta$ Cat # 900-K95, IL-33 Cat #900-K398, hBD1 Cat #900-K202). Pre-coated ELISAs were from Aviva Systems Biology (Muc7 Cat # OKEH01290, Muc5B Cat # OKEH02841, Muc5AC Cat # OKEH02840, lysozyme Cat # OKCD01349, lactoferrin Cat # OKEH02822). Intracellular cAMP from lysed cells was measured using Amersham Biosciences cAMP Biotrak Enzyme Immunoassay system (GE Healthcare) per the manufacturer's instructions using the non-acteylation protocol (sensitivity of 25-6400 fmol per assay well). Live cell imaging of cAMP dutilized downward green cADDis in a baculovirus modified for mammalian cells (BacMam; Montana Molecular). IL-4, IL-13, and LPS were from Cell Signaling Technologies and prepared with carrier BSA as per the manufacturer's instructions. Poly(I:C), NPY, scrambled NPY, BIBO 3304, VIP, $VIP_{(6-28)}$, [D-p-Cl-$Phe^6$,$Leu^{17}$]-VIP, were from Tocris. $T16A_{inh}$-A01, $CaCC_{inh}$-A01, $CFTR_{inh}$-172, NPPB, 4,4'-diisothiocyanato-2,2'-silbenedisulfonic acid (DIDS), carbachol (CCh) were from Cayman Chemical. Inorganic salts for buffers, TNF$\alpha$, bumetanide, forskolin, 4,4'-dinitrostilbene-2,2'-disulfonic acid (DNDS), 5-(N,N-dimethyl)amiloride (DMA), *N*-phenyl-1-naphthylamine (NPN), and all other reagents were from Sigma-Aldrich, unless otherwise indicated below.

Antibodies for lysozyme (BGN/06/96I; Cat # ab36362), lactoferrin (2B8, Cat # ab10110), $Na^+/K^+$ ATPase (EP1845Y; Cat # ab76020), Muc7 (Cat # ab55542), Muc5AC (Cat # ab3649), alpha-1-antitrypsin (Cat # ab20830), $\beta$2 adrenergic receptor (Cat # ab182136) and Glut1 (rabbit polyclonal; Cat# ab15039) were from Abcam. Mouse monoclonal antibody to Glut1 (SPM498; Cat # MS-10637) was from Thermo. Antibodies to VIPR1 (Cat # AVR-001), TMEM16A (Cat # ACL-011), CFTR (Cat # ACL-006) were from Alomone Labs. Antibody to VIPR2 (Cat # PA3-114) was from Pierce.

All solutions used were prepared as described (1, 2). Krebs $HCO_3^-$ buffer for isolated acinar cell experiments contained 125 NaCl, 5 KCl, 1.2 $MgCl_2$, 1.2 $NaH_2PO_4$, 11 glucose, and 25 $NaHCO_3$, gassed with 95% $O_2$ + 5% $CO_2$. Krebs $HCO_3^-$ -free buffer contained 125 NaCl, 5 KCl, 1.2 $MgCl_2$, 1.2 $CaCl_2$, 1.2 $NaH_2PO_4$, 11 glucose, 20 Hepes, 20 sucrose, pH 7.4, gassed with 100% $O_2$. Solutions for buffering capacity measurement, SNARF calibration, and SPQ calibration were as described (1, 2) and are indicated below. Hank's balanced salt solution (HBSS) contained (in mM) 138 NaCl, 5.3 KCl, 0.4 $KH_2PO_4$, 0.34 $NaHPO_4$, 0.41 $MgSO_4$, 0.49 $MgCl_2$, 1.8 $CaCl_2$, 5.6 glucose, 20 mM HEPES pH 7.4. Unless indicated, all cell culture reagents were from Gibco.

***Isolation and culture of primary nasal gland serous acinar cells***

Isolated tissue was first placed in HBSS supplemented 2 mm L-glutamine, MEM-vitamins MEM-amino acids, MEM non-essential amino acids, and 1% BSA. The epithelium was removed via forceps and submucosal tissue was removed from the bone. The tissue was mechanically minced with scissors and then incubated for 90 min at room temperature in HBSS supplemented as above but with 1 mg/ml Collagenase P (Roche) and 10 µg/ml DNAse I (Roche) with gentle shaking. Remaining intact tissue pieces were separated from dispersed acini and acinar cells by gravity (3 min). Gland acini were separated from single epithelial or immune cells by a short centrifugation (30 sec, 500x g). The isolation protocol yielded acini and strings of acinar cells. Acini were further dispersed by incubation with 0.5 mg/ml collagenase P as above for 60 min. Cells were pelleted and washed with HBSS before being seeded onto glass coverslips for imaging or collagen-coated transwells.

***Immunofluorescence (IF)***

IF was carried out as previously described (3), with modifications outlined below. ALI cultures were fixed for 3 min in ice-cold methanol, followed by blocking in Dulbecco's phosphate buffered saline (DPBS) containing 1% bovine serum albumin (BSA), 5% normal donkey serum (NDS), 0.2% saponin, and 0.3% triton X-100 for 1 hour at 4°C. Primary antibody incubation was carried out at 4°C overnight. AlexaFluor-labeled donkey anti-mouse or rabbit secondary antibody incubation (1:1000) was carried out for 2 hours at 4°C. Transwell filters were removed from the plastic mounting ring and mounted with Fluoroshield with DAPI (Abcam). Images of ALIs were taken on an Olympus IX83 microscopy (60x 1.4 NA objective) with spinning disc confocal unit (Olympus DSU). Images were analyzed using Metamorph software and/or the FIJI (4) version of ImageJ (W. Rasband, Research Services Branch, National Institute of Mental Health, Bethesda, MD).

***SNARF-5F, SPQ, and DIC live-cell imaging of primary isolated serous cells***

Isolation of primary serous acinar cells, immunofluorescence, and live cell imaging of acinar cell volume, pHi (SNARF-5F), and $Cl^-$ (SPQ) was carried out as described (1, 2, 5-7). After washing via gentle centrifugation and resuspension in $HCO_3^-$ containing buffer, acinar cells were plated on Cell-Tak (BD Biosciences)-coated glass coverslips and allowed to adhere for 10–20 min in 5% $CO_2$. The isolation protocol yielded acini, single cells and strings of cells. Cells were identified based on visible morphology (size, polarized secretory granules, acinar structures) under DIC optics.

Isolated acinar cells were loaded with SNARF-5F-acetoxymethyl ester (AM) for 15 min at room temperature in Krebs buffer containing 25 mM $HCO_3^-$ gassed with 5% $CO_2$/95% $O_2$. For experiments in the absence of $HCO_3^-$, cells were plated in Krebs buffer lacking $NaHCO_3$ but containing 20 mM HEPES and gassed with 100% $O_2$. Solutions with 0-$Na^+$ had isosmotic replacement of NaCl with NMDG-Cl, $NaH_2PO_4$ with $KH_2PO_4$, and $NaHCO_3$ with NMDG-$HCO_3$. Solutions used were exactly as previously described (2). Ratiometric fluorescence measurements of SNARF-5F were carried out using 550/20 excitation filter, 570 long pass dichroic, and 585/20 and 640/20 emission filters (Chroma Technologies set 79010-ET) housed in a filter wheel (Sutter). Excitation light was generated with an X-Cite 120 Boost LED (Excelitas Technologies) and emission was captured with an ORCA Flash 4.0 sCMOS camera (Hamamatsu) with 2x2 pixel binning. Imaging was performed on a Olympus IX-83 microscope with 30x 1.05 NA UPlanSApo silicone oil immersion objective for single cell measurements or 10x 0.4 NA PlanApo lens for ASL measurements with cells on transwells. Single cells were continuously perfused with 37°C solution gassed with 95% $O_2$/5% $CO_2$ or 100% $O_2$ as appropriate. Transwells were kept at 37°C with 5% $CO_2$ using a Tokai Hit stage-top incubator.

For experiments blocking the driving force for $HCO_3^-$ and $Cl^-$ efflux (performed as described (2), we assumed $[HCO_3^-]_i$ = 16 mM based on mean resting pH of 7.2 and $[HCO_3^-]_o$ = 25 mM, making the Nernst equilibrium potential ($E_{HCO3-}$) ~60mV•log(16/25) = -12 mV. We have already demonstrated that the activation of secretion by serous cells results in efflux of KCl (1, 2, 6, 7). Mean resting $[Cl^-]_i$ was measured at ~65 mM in SPQ experiments, and with $[Cl^-]_o$ in the Krebs buffer used here at 135 mM, for a $E_{Cl-}$ = -19 mV. $[K^+]_i$ was assumed to be 140 mM and $[K^+]_o$ was calculated at 5 mM ($E_{K+}$ = -87 mV). Using the Nernst equation, we calculated that a using $[Cl^-]_o$ of 103 mM and $[K^+]_o$ of 89 mM would set $E_{Cl-} = E_{K+} = E_{HCO3-}$, reducing the driving force for efflux of cellular KCl and $KHCO_3$. This solution contained (in mM) 41 NaCl, 57 KCl, 32 KGluconate, 1.2 $MgCl_2$, 1 $CaCl_2$, 1.2 $NaH_2PO_4$, 11 glucose, 25 $NaHCO_3$, pH 7.4 by gassing with 95% $O_2$/5% $CO_2$ compared with control Krebs that contained (in mM) 125 NaCl, 5 KCl, 1.2 $MgCl_2$, 1.2 $CaCl_2$, 1.2 $NaH_2PO_4$, 11 glucose, 25 $NaHCO_3$, pH 7.4 by gassing with 95% $O_2$/5% $CO_2$.

SPQ measurement of $[Cl^-]_i$ changes were carried out exactly as described (1, 2, 5-8). Isolated acinar cells were incubated for 2 hours in 20 mM SPQ at room temperaure. Acinar cell ALIs were incubated overnight with 20 mM SPQ on the apical side. SPQ was imaged using a standard DAPI filter set (350/50 ex, 400 long pass dichroic, 460/50 em; Chroma 49000 ET) with UV illumination from a xenon arc lamp (Sutter Lamda LS). Solutions used for $NO_3^-$ substitution were as previously described (1, 5-8). For ALI experiments, control normal $[Cl^-]_o$ apical solution contained (in mM) 138 NaCl, 5.3 KCl, 0.24 $MgCl_2$, 1.3 $CaCl_2$ (total $[Cl^-]_o$ = 147), 20 HEPES pH 7.4. Low $[Cl^-]_o$ solution contained (in mM) 138 $NaNO_3$ and 5.3 $KNO_3$ instead of NaCl and KCl, respectively (final $[Cl^-]_o$ = 4; ~37-fold less than normal $[Cl^-]_o$). For isolated acinar cells, control solution contained (in mM), 136.2 NaCl, 3.8 KCl, 1.2 KH2PO4, 1.2 CaCl2, 1.2 MgCl2, 11 glucose, 10 HEPES pH 7.4. Low $[Cl^-]_o$ solution contained NaCl replaced with $NaNO_3$ for a final $[Cl^-]_o$ of 8.6 mM. Single cells were continuously perfused with 37°C solution gassed with 95% $O_2$/5% $CO_2$ or 100% $O_2$ as appropriate. Transwell SPQ experiments were carried out at room temperature without gassing.

Cell volume was estimated by taking the cross-sectional area of the cell as imaged by differential interference contrast (DIC) to the 3/2 power (as described (1, 5-7, 9-11)). This method yields cell volume measurements faster but indistinguishable from confocal 3D reconstructions (1). Cell volumes are expressed as normalized volume (V) relative to initial cell volume ($V_o$). DIC images were acquired sequentially by computer controlled shuttering off of the fluorescence light, rotating of the DIC polarizer into position, and shuttering on transmitted light. Imaging data was collected and analyzed in Metafluor and/or FIJI (4).

***Calibration of SNARF-5F and measurement of intracellular pH ($pH_i$) buffering capacity***

Changes in SNARF 640/580 emission ratio were converted to pHi using SNARF-loaded cells exposed high $[K^+]_o$ solutions and the $H^+/K^+$ exchanger nigericin to equilibrate extracellular pH ($pH_o$) to $pH_i$ exactly as described (2) using solutions buffered to $pH_o$ 6.8, 7.2, and 7.6. SNARF-5F fluorescence varied linearly within the pH ranges observed during agonist stimulation.

Total $pH_i$ buffering capacity ($\beta_t$) encompasses $CO_2$-$HCO_3^-$ -dependent buffering capacity ($\beta_{HCO3-}$) plus intrinsic $CO_2$-independent intrinsic buffering capacity ($\beta_i$) from cytoplasmic macromolecules and organelles (2, 12, 13). Assuming the pKa of $CO_2$-$HCO_3^-$ is 6.1 (2, 13, 14) and assuming that highly permeant $[CO_2]_o = [CO_2]_i$ (1.2 mM in 5% $CO_2$ by Henry's Law), and using the Henderson-Hasselbach relationship, then $[HCO_3^-]_i = 1.2$ mM x $10^{pH-6.1}$, and $\beta_{HCO3-} = 2.3 \times [HCO_3^-]_i$. Because $[CO_2]_i$ is constant (open buffering), $\beta_{HCO3-}$ rises exponentially as $pH_i$ increases (2).

Human serous acinar cell $\beta_i$ was empirically determined using observed $pH_i$ changes during exposure to $NH_4Cl$ in $Na^+/HCO_3^-$ -free solutions to inhibit $pH_i$ regulatory mechanisms (as described; (2, 12, 13, 15)). Exposure of cells to solution containing $NH_3$ and $NH_4^+$ causes an initial alkalinization of $pH_i$ due to entry of highly cell permeant $NH_3$ and resulting $H^+$ consumption as it is converted intracellularly to $NH_4^+$. After an experimental change in extracellular [$NH_3$] ([$NH_3$]$_o$), the initial intracellular [$NH_4^+$]$_i$ can be calculated by Henderson-Hasselbach with [$NH_4^+$]$_i$ = [$NH_3$]$_i$ x $10^{9.2\text{-pHi}}$, assuming [$NH_3$]$_o$ = [$NH_3$]$_i$ are identical and pKa = 9.2 (13). Acinar cells were exposed to solutions containing (in mM) 0, 5, 10, and 20 mM [$NH_4Cl$]$_o$, which contained (in mM) 0, 0.6, 1.2, and 2.5 [$NH_3$]$_o$. The base solution for $\beta_i$ buffering experiments was (in mM) 120-140 NDMG-Cl, 5 KCl, 1.2 $MgCl_2$, 1.2 $CaCl_2$, 1.2 $KH_2PO_4$, 11 glucose, 10 HEPES pH 7.4, and 0, 5, 10 or 20 $NH_4Cl$ gassed with 100% $O_2$. Mean $\beta_i$ was calculated as the units of acid of base equivalent required to change the $pH_i$ by one unit around the midpoint of the pH change as described (2). Raw data points for $\beta_i$ were taken from experiments of 12 cells of each genotype (4 patients; 3 experiments per patient) and fit with an exponential decay function in Prism. The sum of the $\beta_i$ and $\beta_{HCO3-}$ curves was used to calculate $\beta_t$.

***Measurements of ASL pH and ASL height***

ASL height and pH was carried out as described (3, 8, 16-22). Cultures were imaged at 37°C in a Tokai Hit stage top incubator. For pH measurements, cells were incubated in serum-free phenol-red-free low glucose DMEM (Gibco) on the basolateral side and gassed with 5% $CO_2$, 20% $CO_2$, 80% N2. For “thin film” ASL pH measurements (main text) SNARF-1 dextran (~1 mg/ml) was sonicated in perfluorocarbon and 100 uL was added to the top of each culture. For longer-term $HCO_3^-$ secretion experiments (**Supplemental Figure 8**), 100 uL of 1 mg/ml SNARF dextran in low buffering capacity solution was added (HBSS with 1 mM HEPES, as described (22)). ASL pH was calibrated by overlaying 1 mg/ml SNARF dextran on top of cultures in $HCO_3^-$ conditions in solutions buffered with 20 mM HEPES at pH 6.8, 7.2, 7.6, and 7.8. SNARF 1 dextran pH changes were linear over the pH range observed here (~7-7.8).

ASL height was measured similarly and as previously described (19), but in $HCO_3^-$ -free conditions (100% $O_2$ with basolateral HBSS buffered with 20 mM HEPES) using Texas red dextran (10,000 MW) as previously described (19). When corrected for refractive index mismatch (1.52

$\eta_{oil}/1.33\ \eta_{water}$ = ~1.14), an observed change in ASL height of ~30 µm with agonist stimulation is in reality 30/1.14 = 26 µm. Treating the ALI as a cylinder, where volume = area x height, a change in ASL height of ~26 µm over 15 min equals a secretion volume of 2.6 µL/cm$^2$ (2.6x10$^{-5}$ m x 1x10$^{-4}$ m$^2$ = 2.6 x 10$^{-9}$ m$^3$ = 2.6x10$^{-6}$ L) or ~10 µL/cm$^2$/hour. Calu-3 cells were previously reported to secrete fluid at a rate of 4 or 5.4 uL•cm$^2$•hr when stimulated with forskolin or VIP, respectively, using a virtual gland technique (23). The fact that measurements of primary serous cells here using the Texas red ASL technique are within an order of magnitude of measurements of Calu-3 cells using a different technique suggests the ASL height measurements made here are reasonable within the context of cellular fluid secretion capabilities.

***Generation of Calu-3 air-liquid interface (ALI) cultures***

Calu-3 bronchial epithelial cells were obtained from ATCC and cultured in T75 flasks in minimal essential medium (MEM) with Earl's salts and 1 mM L-glutamine, 10% fetal bovine serum, and 1% penicillin/streptomycin mix. Cells were lifted with 0.25% trypsin and plated on 1.1 cm$^2$ cell culture inserts (Greiner BioOne Thincerts, transparent, 0.4 µm pore size). Cells were grown to confluence for 5 days, followed by apical exposure to air and subsequent 3 weeks for full differentiation/polarization before use. Only ALIs with transepithelial resistances (TEERs) of >250-300 Ω•cm$^2$ were used.

***Bacterial growth assays***

Bacterial growth assays were carried out as previously described (24, 25). *Pseudomonas aeruginosa* strains PAO1 (HER-1018; ATCC BAA-47) and clinical isolates of methicillin-resistant *Staphylococcus aureus* (MRSA) and *P. aeruginosa* were isolated by the Philadelphia VA Medical Center Microbiology Laboratory and grown in LB or tryptic soy broth (TSB; Gibco/Thermo Scientific), respectively.

Bacterial NPN fluorescence assay was modified from previous descriptions (26-29). *P. aeruginosa* were grown to an $OD_{600}$ of 0.5 in LB, centrifuged, and resuspended at half volume of 10 mM HEPES, 5 mM glucose, 0.1 mM EDTA, pH 8. Bacteria were then aliquoted and mixed with an equal volume of diluted airway surface liquid secretions or antibiotics, and then pipetted into a plate reader containing an equal volume of 25% PBS containing 20 µM NPN (final NPN 10 µM, final $OD_{600}$ 0.25). Samples were then incubated for 10 min and read on a Tecan 10M plate reader at 350 nm excitation and 450 nm emission. Samples were read in triplicate, with averages of at least 3 independent experiments reported.

CFU antimicrobial assays with Calu-3 ASL washings were carried out similarly to a previously published protocol (18, 30) and modified based on our own antimicrobial ASL protocols used in our lab (18). Cultures were washed copiously with PBS and transferred to antibiotic-free MEME for 48 hrs. before use. Calu-3 cell secretions were collected from 3 week old ALIs stimulated basolaterally with 100 µM isoproterenol for 72 hours, followed by washing of the apical surface with 30 µL 25% PBS. While washing a 1.1 $cm^2$ ALIs with 30 µL significantly dilutes the ASL fluid (~1 µL per $cm^2$ of surface area (31)), washings retained antibacterial activity and were thus sufficient to be used for this assay. ASL washings (30 µL per culture) were pooled and mixed with bacteria resuspended in 25% PBS, adjusted to 0.1 OD, then diluted 1:1000 in 25% PBS). Bacteria and ASL mixture was incubated statically in a 96-well plate at 37 °C for 2 hrs, followed by 4 serial 10-fold dilutions and spot plating onto LB plates. After overnight incubation at 37 °C, CFUs were manually counted.

CFU antimicrobial assays with primary serous cell ASL washings were carried out as above, but cultures were not pre-treated with isoproterenol. Cultures were unstimulated or stimulated for 30 min with VIP ± NPY ± scrambled NPY on the basolateral side. Afterward, the surface of a 0.33 $cm^2$ transwell was washed with 50 uL 25% saline (thus ASL was ~5x more dilute than used in Calu-3 experiments).

Live-dead staining was carried out with BacLight Live/Dead kit (ThermoFisher Scientific) consisting of Syto9 (live cell stain) and propidium iodide (dead cell stain), as previously used with *P.*

*aeruginosa* (3). Bacteria were adjusted to an OD = 0.1. Bacterial suspension was mixed with ASL (25 µL each) in a black microplate and incubated for indicated time at 37 °C. 50 µL 2x Live-Dead staining solution was then added, followed by further 10 min incubation at room temp and reading on a fluorescence microplate reader (Tecan Spark 10M) at 488 excitation and dual emission wavelengths as indicated. Control calibration of live dead staining was carried out by mixing heat-killed (as below) with live bacteria at the indicated ratios to a final OD of 0.1 followed by mixing with 25 % saline only and incubation as above prior to live dead staining.

***Production of heat-killed bacteria***

Bacteria were heat killed according to a previously published protocol (32). *P. aeruginosa* or MRSA strains were grown overnight at 37°C in LB broth, then resuspended in LB and grown for 2-4 hours to an OD600 of 1. Bacteria were heat killed for 20 min at 95°C. Cells were treated with bacteria diluted to $OD_{600}$ = 0.01 (100x) in 100 uL PBS on the apical side only. Unstimulated control cultures were treated with 1:100 LB media only.

**Supplemental Table 1.** Gene expression output from the Cancer Cell Line Atlas (accessed 26 April, 2019; https://portals.broadinstitute.org/ccle) for NPY receptors, VIP receptors, and serous cell markers lysozyme (LYZ) and CFTR. Note that Calu-3 cells, a bronchial adenocarcinoma line frequently used as a model of serous cells due to high CFTR and lysozyme expression, express the highest amount of NPY1R relative to other airway cancer cell lines.

Affymetrix

| Gene | NPY1R | NPY2R | NPY5R | VIPR1 | VIPR2 | LYZ | CFTR |
|---|---|---|---|---|---|---|---|
| A549_LUNG | 3.97359 | 3.910246 | 4.316899 | 5.347613 | 4.442829 | 3.861464 | 3.98063 |
| CALU1_LUNG | 4.150831 | 3.587063 | 4.239859 | 7.542943 | 4.087702 | 3.718109 | 3.854273 |
| CALU3_LUNG | 7.168295 | 3.803298 | 4.45895 | 6.922686 | 4.311346 | 11.0784 | 10.05969 |
| CALU6_LUNG | 3.512282 | 3.750993 | 4.158273 | 8.829339 | 4.305164 | 6.284744 | 4.588329 |
| NCIH292_LUNG | 3.553423 | 3.486564 | 3.898884 | 6.090106 | 4.314193 | 3.713948 | 3.729571 |
| NCIH441_LUNG | 3.701913 | 3.58951 | 4.291047 | 7.25233 | 4.440556 | 3.726355 | 4.128871 |
| NCIH520_LUNG | 3.952049 | 3.850606 | 4.148694 | 5.315201 | 4.260294 | 4.190855 | 4.465682 |
| NCIH522_LUNG | 3.666311 | 3.629025 | 4.243514 | 5.523483 | 4.433325 | 3.896822 | 3.716021 |

RNAseq

| Gene | NPY1R | NPY2R | NPY4R | NPY5R | VIPR1 | VIPR2 | LYZ | CFTR |
|---|---|---|---|---|---|---|---|---|
| A549_LUNG | -5.3022356 | -3.1772462 | 5.69438127 | -13 | -4.4839539 | -8.4031139 | -4.837362 | -6.4493319 |
| CALU1_LUNG | -1.6370841 | -13 | -0.0247348 | -13 | 1.83239394 | -13 | -3.4844883 | -4.3859648 |
| CALU3_LUNG | 0.76601676 | -13 | -2.4547363 | -1.7320273 | 0.81640217 | -13 | 7.36181905 | 6.44949113 |
| CALU6_LUNG | -4.1292877 | -7.6433373 | -5.1671067 | -6.7017855 | 3.62349708 | -9.1687653 | 1.71665862 | 0.66432141 |
| NCIH292_LUNG | -8.8698593 | -13 | 0.67477717 | -13 | 0.85130002 | -0.6510655 | -2.2350607 | -7.3064622 |
| NCIH441_LUNG | -13 | -13 | -4.4713423 | -13 | 1.79193401 | -6.2665501 | -3.3788701 | -5.7721996 |
| NCIH520_LUNG | -4.5970396 | -13 | -6.0181873 | -13 | -3.6856484 | -9.4348835 | -0.494092 | -3.896139 |
| NCIH522_LUNG | -7.686093 | -13 | -6.7853125 | -13 | -3.0377362 | -6.6170462 | -1.6362569 | -5.7076583 |

**Supplemental Table 2.** Gene expression output from MEERAV database (http://meerav.wi.mit.edu). Similarly to the Cancer Cell Line Atlas data above, MEERAV (accessed 11 July 2018) suggested Calu-3 cells expressed the highest amount of NPY1R among frequently used lung cancer cell lines.

| GENE | NPY1R | NPY2R | NPY4R | NPY5R | NPY6R | VIPR1 | VIPR2 | LYZ | CFTR |
|---|---|---|---|---|---|---|---|---|---|
| A549 Lung_Cancer.Cell.Line_Large_A549_Non-Small.Cell.Lung.Carcinoma: METIS_p_NCLE_RNA1_Human_U133_Plus_2.0_H01_241018 | 20.28 | 19.39 | 104.89 | 29.99 | 64.41 | 69.52 | 54 | 28.38 | 24.6 |
| A549 Lung_Cancer.Cell.Line_Large_A549_Non-Small.Cell.Lung.Carcinoma: GSM139646 | 16.49 | 17.66 | 78.19 | 20.19 | 66.14 | 73.3 | 58.19 | 26.4 | 24.02 |
| A549 Lung_Cancer.Cell.Line_Large_A549_Non-Small.Cell.Lung.Carcinoma: GSM253203 | 26.74 | 19.48 | 46.55 | 43.68 | 136.8 | 130.62 | 54.08 | 24.67 | 24.66 |
| A549 Lung_Cancer.Cell.Line_Large_A549_Non-Small.Cell.Lung.Carcinoma: GSM274739 | 18.43 | 17.99 | 111.42 | 36.73 | 87.56 | 81.09 | 50.29 | 25.34 | 24.33 |
| A549 Lung_Cancer.Cell.Line_Large_A549_Non-Small.Cell.Lung.Carcinoma: GSM274740 | 20.94 | 20.97 | 141.78 | 35.76 | 83.06 | 77.51 | 42.31 | 24.14 | 24.59 |
| A549 Lung_Cancer.Cell.Line_Large_A549_Non-Small.Cell.Lung.Carcinoma: GSM433024 | 55.74 | 20.57 | 76.52 | 39.55 | 70.34 | 95.51 | 55.45 | 25.13 | 26.5 |
| A549 Lung_Cancer.Cell.Line_Large_A549_Non-Small.Cell.Lung.Carcinoma: GSM433025 | 57.1 | 24.58 | 81.78 | 31.01 | 66.64 | 92.39 | 52.3 | 28.06 | 20.06 |
| Calu3 Lung_Cancer.Cell.Line_Large_Calu-3_Adenocarcinoma: WATCH_p_NCLE_RNA8_HG-U133_Plus_2_A02_474612 | 128.77 | 18 | 45.06 | 34.25 | 66.32 | 195.12 | 47.65 | 872.07 | 708.73 |
| Calu6 Lung_Cancer.Cell.Line_Large_Calu-6_Adenocarcinoma: BRAKE_p_NCLE_RNA2_HG-U133_Plus_2_E04_241144 | 13.41 | 16.41 | 50.21 | 26.83 | 94.21 | 447.11 | 44.95 | 68.77 | 32.67 |
| Calu 6 Lung_Cancer.Cell.Line_Large_Calu-6_Adenocarcinoma: Calu-6_SS474757_HG-U133_Plus_2_HCHP-225882_ | 17.07 | 15.34 | 32.14 | 40.08 | 73.67 | 564.18 | 45.56 | 105.84 | 26.02 |
| Calu-6 Lung_Cancer.Cell.Line_Large_Calu-6_Adenocarcinoma: Calu-6_SS474758_HG-U133_Plus_2_HCHP-225883_ | 16.23 | 17.07 | 30.89 | 36.32 | 79.62 | 567.31 | 44.38 | 109.59 | 25.33 |
| Calu-6 Lung_Cancer.Cell.Line_Large_Calu-6_Adenocarcinoma: Calu-6_SS474759_HG-U133_Plus_2_HCHP-225884_ | 19.51 | 17.4 | 34.71 | 32.74 | 69.48 | 552.72 | 47.16 | 104.84 | 25.04 |
| H292 Lung_Cancer.Cell.Line_Large_NCI-H292_Mucoepidermoid.Carcinoma: GSM274745 | 15.82 | 14.98 | 44.11 | 39.69 | 75.71 | 116.55 | 48.99 | 27.7 | 21.59 |
| H292 Lung_Cancer.Cell.Line_Large_NCI-H292_Mucoepidermoid.Carcinoma: GSM274746 | 15.69 | 16.33 | 47.04 | 34.49 | 74.92 | 89.52 | 50.44 | 24.76 | 22.58 |
| H441 Lung_Cancer.Cell.Line_Large_NCI-H441_Bronchioloalveolar.Adenocarcinoma: METIS_p_NCLE_RNA1_Human_U133_Plus_2.0_E05_240954 | 14.87 | 16.62 | 49.04 | 34.08 | 87.71 | 212.35 | 48.29 | 18.77 | 24.56 |
| H441 Lung_Cancer.Cell.Line_Large_NCI-H441_Bronchioloalveolar.Adenocarcinoma: GSM274741 | 16.59 | 14.5 | 38.25 | 45.12 | 72.27 | 119.51 | 47.77 | 26.89 | 22.77 |
| H441 Lung_Cancer.Cell.Line_Large_NCI-H441_Bronchioloalveolar.Adenocarcinoma: GSM274742 | 14.95 | 15.78 | 44.38 | 37.62 | 71.5 | 116.88 | 44.24 | 25.52 | 21.04 |
| H441 Lung_Cancer.Cell.Line_Large_NCI-H441_Bronchioloalveolar.Adenocarcinoma: NCI-H441_SS475525_HG-U133_Plus_2_HCHP-225837_ | 15.89 | 18.07 | 48.6 | 28.07 | 69.86 | 263.85 | 43.29 | 23.61 | 28.88 |
| H441 Lung_Cancer.Cell.Line_Large_NCI-H441_Bronchioloalveolar.Adenocarcinoma: NCI-H441_SS475527_HG-U133_Plus_2_HCHP-225839_ | 19.1 | 16.67 | 50.57 | 27.93 | 68.59 | 226.22 | 44.21 | 26.03 | 27.01 |
| H520 Lung_Cancer.Cell.Line_Large_NCI-H520_Squamous.Cell.Carcinoma: BRAKE_p_NCLE_RNA2_HG-U133_Plus_2_D02_241116 | 19.37 | 17.66 | 59.98 | 28.5 | 76.65 | 67.04 | 44.04 | 33.57 | 34.41 |
| H520 Lung_Cancer.Cell.Line_Large_NCI-H520_Squamous.Cell.Carcinoma: GSM274798 | 16.5 | 15.8 | 48.05 | 30.93 | 78.98 | 87.21 | 46.82 | 36.39 | 21.66 |
| H520 Lung_Cancer.Cell.Line_Large_NCI-H520_Squamous.Cell.Carcinoma: GSM274831 | 20.61 | 16.66 | 47.85 | 34.97 | 77.36 | 82.98 | 45.82 | 32.57 | 25.09 |
| H522 Lung_Cancer.Cell.Line_Large_NCI-H522_Adenocarcinoma: NIECE_p_NCLE_RNA3_HG-U133_Plus_2_B08_296028 | 14.91 | 17.14 | 42.85 | 31.33 | 98.42 | 94.97 | 57.63 | 24.73 | 21.26 |
| H522 Lung_Cancer.Cell.Line_Large_NCI-H522_Adenocarcinoma: GSM274753 | 17.07 | 17.13 | 48.65 | 38.8 | 86.16 | 65.48 | 48.6 | 28.98 | 23.56 |
| H522 Lung_Cancer.Cell.Line_Large_NCI-H522_Adenocarcinoma: GSM274754 | 16.1 | 15.3 | 42.23 | 36.5 | 81.83 | 76.29 | 53.99 | 31.59 | 21.85 |
| H522 Lung_Cancer.Cell.Line_Large_NCI-H522_Non-Small.Cell.Lung.Carcinoma: NCI-H522_SS181856_HG-U133_Plus_2_HCHP-167900_ | 14.4 | 16.48 | 43.98 | 22.48 | 68.53 | 77.74 | 70.22 | 20.51 | 22.26 |
| H522 Lung_Cancer.Cell.Line_Large_NCI-H522_Non-Small.Cell.Lung.Carcinoma: NCI-H522_SS181857_HG-U133_Plus_2_HCHP-167901_ | 15.77 | 16.37 | 39.13 | 22.04 | 110.03 | 67.85 | 54.42 | 22.81 | 23.76 |

#### Supplemental Figure 1

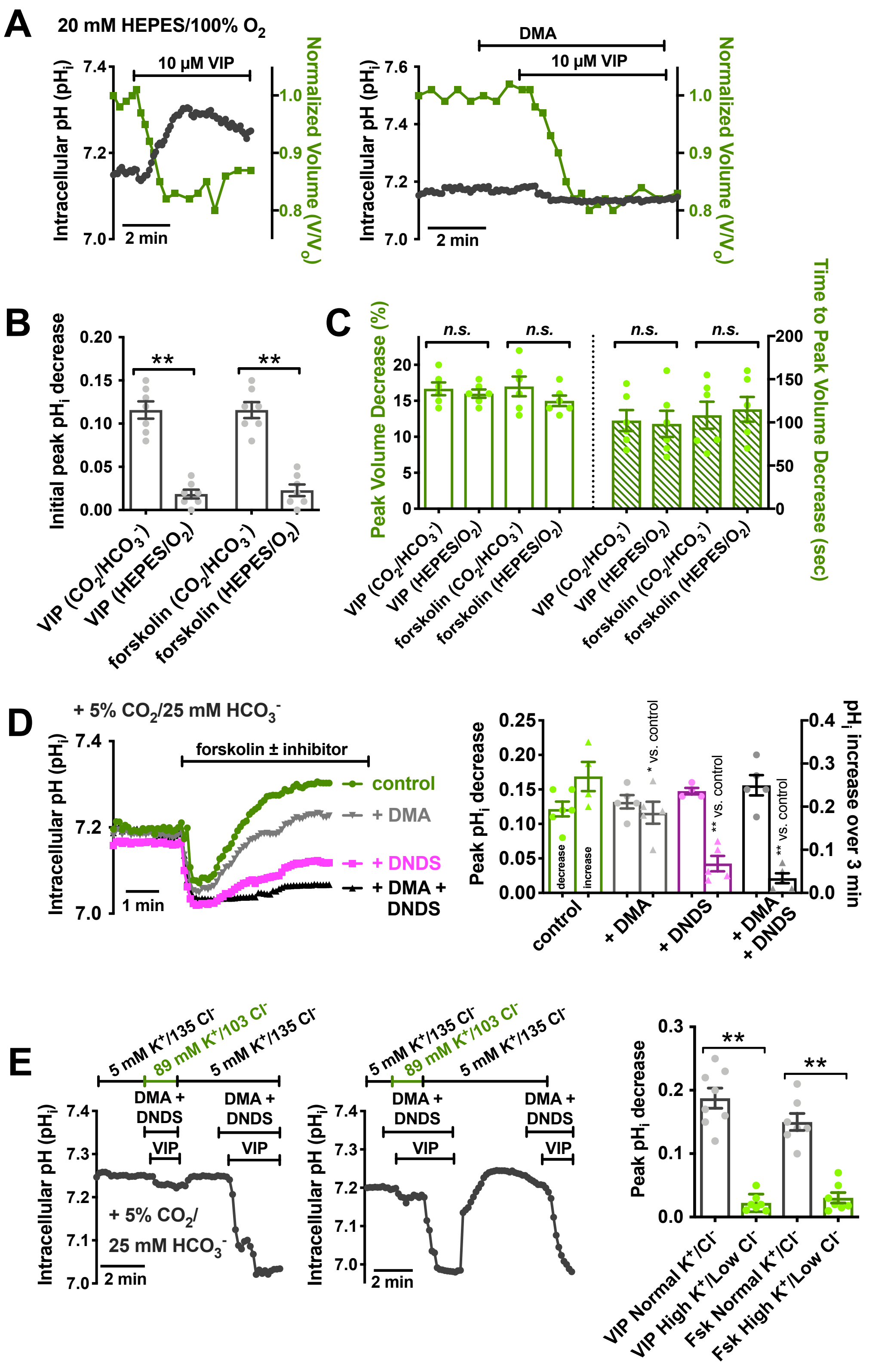

**Supplemental Figure 1: VIP-induced acidification reflects conductive $HCO_3^-$ efflux, while subsequent alkalinization reflects HCO3- uptake via $Na^+HCO_3^-$ cotransporter (NBC). (A).** In the absence of $HCO_3^-$ (20 mM HEPES-buffered conditions gassed with 100% $O_2$), VIP-induced acidification is eliminated. However, cells still shrink at a normal magnitude and rate. Residual $pH_i$ increase is blocked by DMA, suggesting it reflects NHE activity. **(B-C).** Bar graphs (mean ± SEM) showing peak $pH_i$ decrease during VIP or forskolin stimulation in the presence or absence of $CO_2/HCO_3^-$ (*B*) or volume decrease magnitude and kinetics (*C*). Bar graphs show mean ± SEM with significance determined by one-way ANOVA with Bonferroni posttest; ** $p < 0.01$ and *n.s.* = no statistical significance. These data demonstrate that VIP-induced $pH_i$ decrease requires $HCO_3^-$, suggesting it reflects $HCO_3^-$ efflux. However, the magnitude of initial cell shrinkage is not $HCO_3^-$ dependent. This likely reflects the magnitude of $Cl^-$ and $HCO_3^-$ solute from the cell content lost during secretion. A serous acinar cell with resting $[Cl^-]_i$ = ~65 mM (**Supplemental Figure 13** and (1)) loses >50% of cellular $Cl^-$ content (>40 meq•$L^{-1}$) (1, 10). However, the actual $HCO_3^-$ content lost from the cell during secretion under control conditions is smaller; a 7.2 to 7.0 $pH_i$ change would drop $[HCO_3^-]_i$ from ~16 mM to 12 mM (calculated via Henderson Hasselbach). Taking into account the cell volume loss (20%), this is a loss of cellular $HCO_3^-$ content of (1 x 16 meq•$L^{-1}$) – (0.8 x 12 meq•$L^{-1}$) = 6.4 meq•$L^-$ $HCO_3^-$. Thus, cell volume is primarily an indicator of $Cl^-$ secretion while $pH_i$ is primarily an indicator of $HCO_3^-$ changes, as previously observed (2, 33). **(D).** In the presence of $HCO_3^-$, acinar cell $pH_i$ increases (after initial decrease) were substantially reduced by NBC inhibitor 4,4'-dinitrostilbene-2,2'-disulfonic acid (DNDS; 100 μM). Alkalinization was not significantly reduced by $Na^+/H^+$ exchanger (NHE) inhibitor dimethyl amiloride (DMA; 30 μM) alone. All experiments done at 37°C in the presence of 5% $CO_2$. Representative traces shown in A. Bar graph in B shows mean ± SEM; ** = p <0.01 by one-way ANOVA with Bonferroni posttest. These data suggest NBC drives serous cell alkalinization, likely as a way to sustain $HCO_3^-$ secretion due the basolateral localization of NBC in exocrine acinar cells (34-38), similar to what was previously observed with NHE sustaining $HCO_3^-$ secretion during cholinergic-evoked secretion (2). By keeping $[HCO_3^-]_i$ elevated, this will increase the driving force for $HCO_3^-$ efflux across the apical membrane through CFTR. **(E).** In the presence of high $K^+$/low $Cl^-$ conditions designed to block conductive $HCO_3^-$ efflux by clamping $E_{K+} = E_{Cl-} = E_{HCO3-} = V_m$ (described in the methods and (2)), VIP-induced acidification is blocked. Bar graph in B shows mean ± SEM with significance (** = p <0.01) determined via Student's *t* test. DMA (30 μM) + DNDS (100 μM) were used to prevent alkalinization. Thus, the VIP-induced $pH_i$ decrease likely reflects conductive $HCO_3^-$ efflux, likely through CFTR and not a $Cl^-/HCO_3^-$ exchanger.

#### Supplemental Figure 2

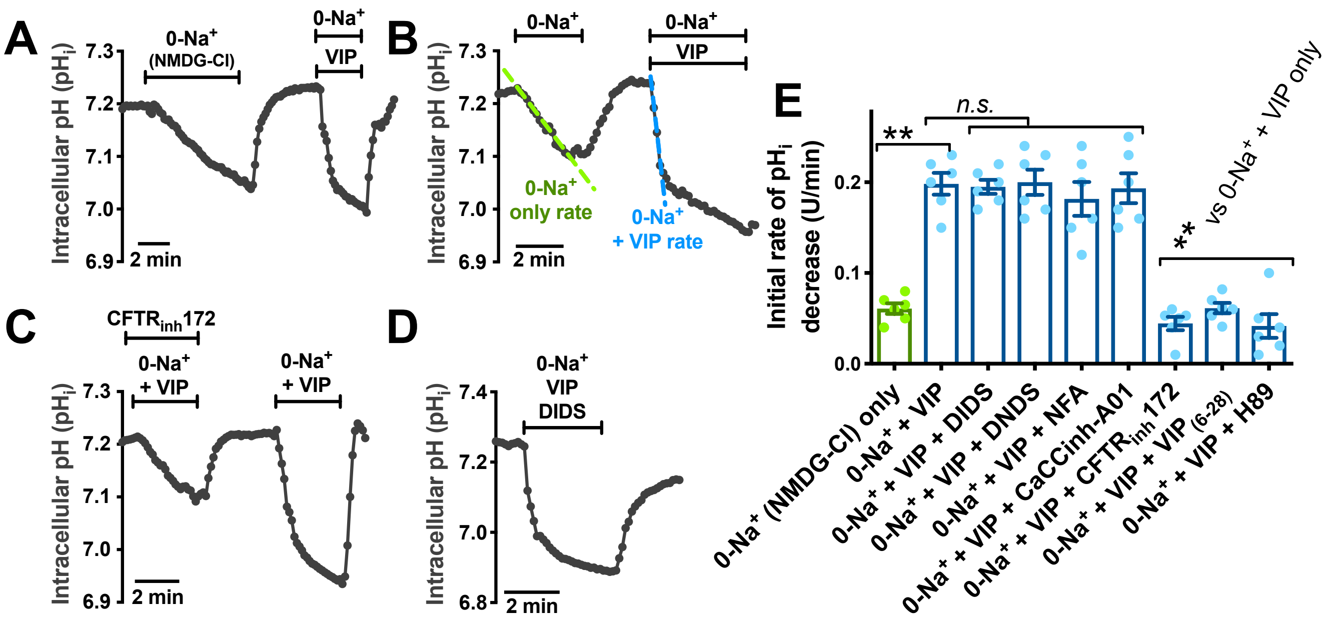


**Supplemental Figure 2: Isolation of the VIP-induced $HCO_3^-$ efflux pathway under 0-$Na^+$ conditions. (A-B).** To better isolate VIP-induced acidification, we performed experiments in 0-$Na^+$ to prevent alkalinization by any $Na^+$ dependent mechanisms (NHE, NBC) in serous cells from non-CF patients. In the absence of $Na^+$ (isosmotic substitution with $NMDG^+$; solutions used described in the Supplemental Methods), cells exhibited a slow acidification. VIP (1 µM) nonetheless still induced an increase in the rate $pH_i$ decrease under these conditions. Panel *B* shows comparisons of rates ± VIP (blue vs green). **(C-D).** The VIP-induced increased in acidification rate was inhibited by $CFTR_{inh}172$ (20 µM; *C*), but not by $Ca^{2+}$-activated $Cl^-$ channel (CaCC) and $Cl^-/HCO_3^-$ exchanger (e.g., pendrin) inhibitors like 4,4'-diisothiocyano-2,2'-stilbenedisulfonic acid (DIDS; *D;* 1 mM), DNDS (30 µM), NFA (100 µM), or $CaCC_{inh}$-A01. **(E).** Bar graphs showing rates measured as in *C-D*. VIP-induced acidification was inhibited only by $CFTR_{inh}172$, VIP receptor antagonist $VIP_{(6-28)}$, or PKA inhibitor H89 (10 µM). Graph shows mean ± SEM with significance determined by one-way ANOVA with Bonferroni posttest; ** $p < 0.01$ and *n.s.* = no statistical significance. All experiments done at 37°C in 5% $CO_2$/25 mM $HCO_3^-$. These data show that inhibitors of TMEM16A/CaCC (39-41) or pendrin (42) do not inhibit VIP-induced acidification. Along with ion substitution (Supplemental Figure 1), these data support the efflux pathway as direct CFTR conduction.

#### Supplemental Figure 3

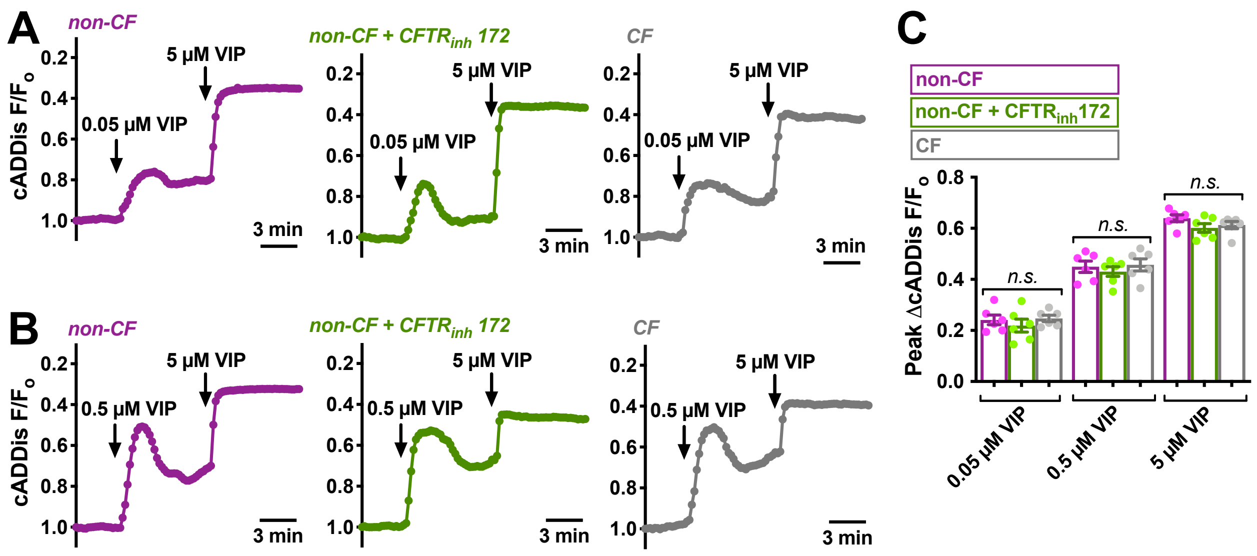


**Supplemental Figure 3: Comparison of VIP-induced cAMP signaling in CF vs non-CF cells.** CFTR has been proposed to act as a hub for kinases and other signaling proteins. We used a fluorescent cAMP biosensor to visualize VIP-activated cAMP increases in CF and non-CF serous acinar cells. Serous acinar cells were isolated, seeded onto CellTak-coated coverslips, and transduced for 6 hrs with a baculovirus pseudotyped for mammalian cells (BacMam) expressing an mNeonGreen-based fluorescent cAMP biosensor (downward cADDis; Montana Molecular, Bozeman MT; (43)) under a CMV promotor followed by 24 hrs incubation. BacMams were previously used to transduce primary lacrimal gland acinar cells (44-46). Single transduced cells and acini were imaged using GFP settings. A decrease in $F/F_o$ (upward deflection of trace) equals an increase in cAMP. **(A-B).** We examined if CF serous cells exhibited alterations in cAMP signaling in response to 0.05, 0.5, and 5 µM VIP. No differences were observed between CF and non-CF patients. This suggests that VIP-evoked cAMP signaling, at least at a global level, is intact in CF serous cells. We also treated non-CF cells with CFTRinh172, and found no alterations of cAMP signals. **(C).** Bar graph of peak responses from representative experiments as shown in *A-B* (3-5 patients for each group, at least 2 experiments per patient per group). 1-way ANOVA with Bonferroni posttest suggested no statistically significant differences. Bar graph shows mean ± SEM; *n.s.* = no statistical significance.

#### Supplemental Figure 4

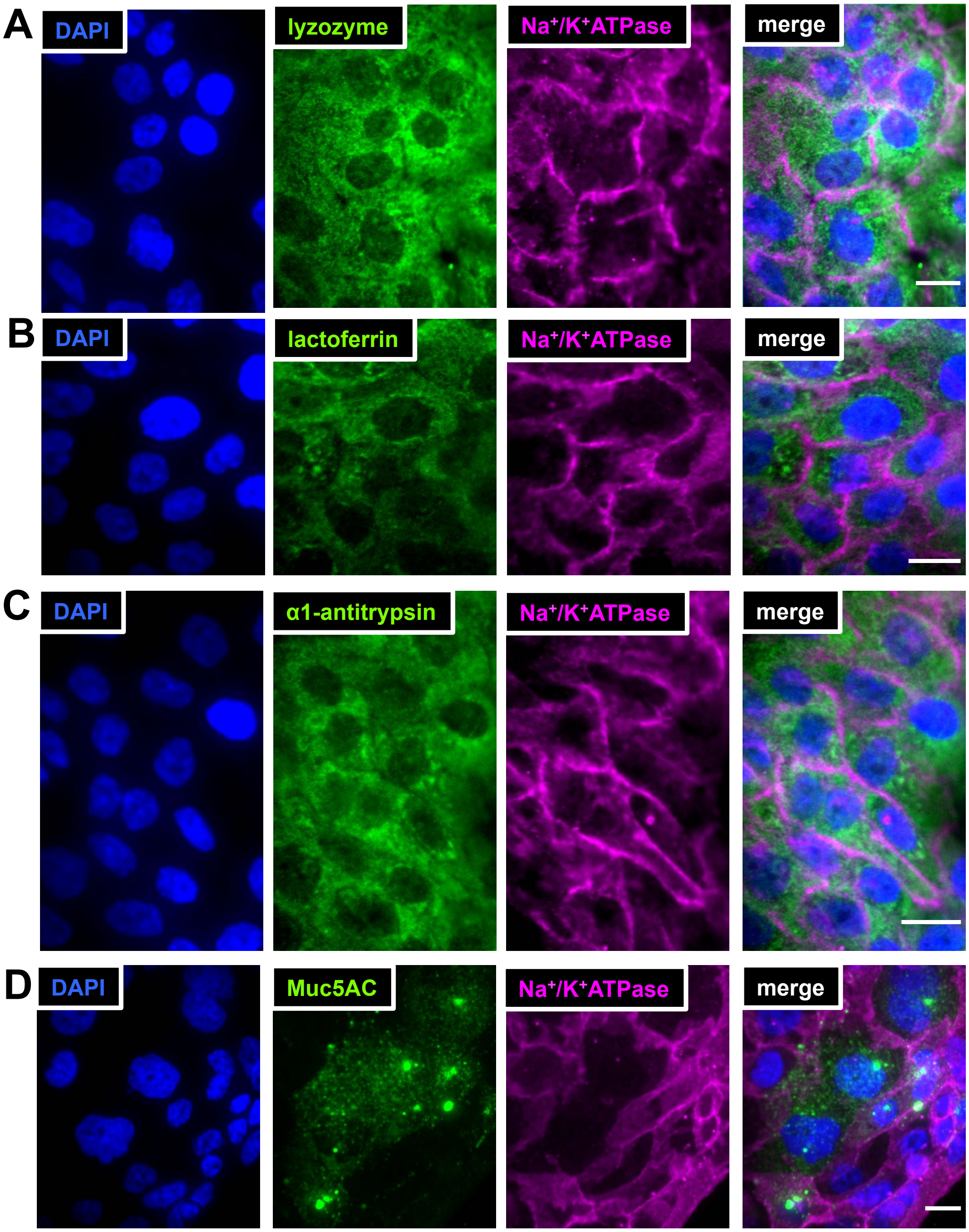


**Supplemental Figure 4: Expression of serous cell markers lysozyme (*A*), lactoferrin (*B*), and antitrypsin (*C*) in Calu-3 cells as well as goblet cell marker Muc5AC (*D*).** Cells were seeded and grown as a monolaryer on collagen coated glass bottom dishes (MatTek), and confluent monolayers were fixed in ice cold MeOH for 3 min before immunostaining as described in the Supplemental Methods. Antibody against $Na^+/K^+$ ATPase was used as a positive plasma membrane control.

#### Supplemental Figure 5

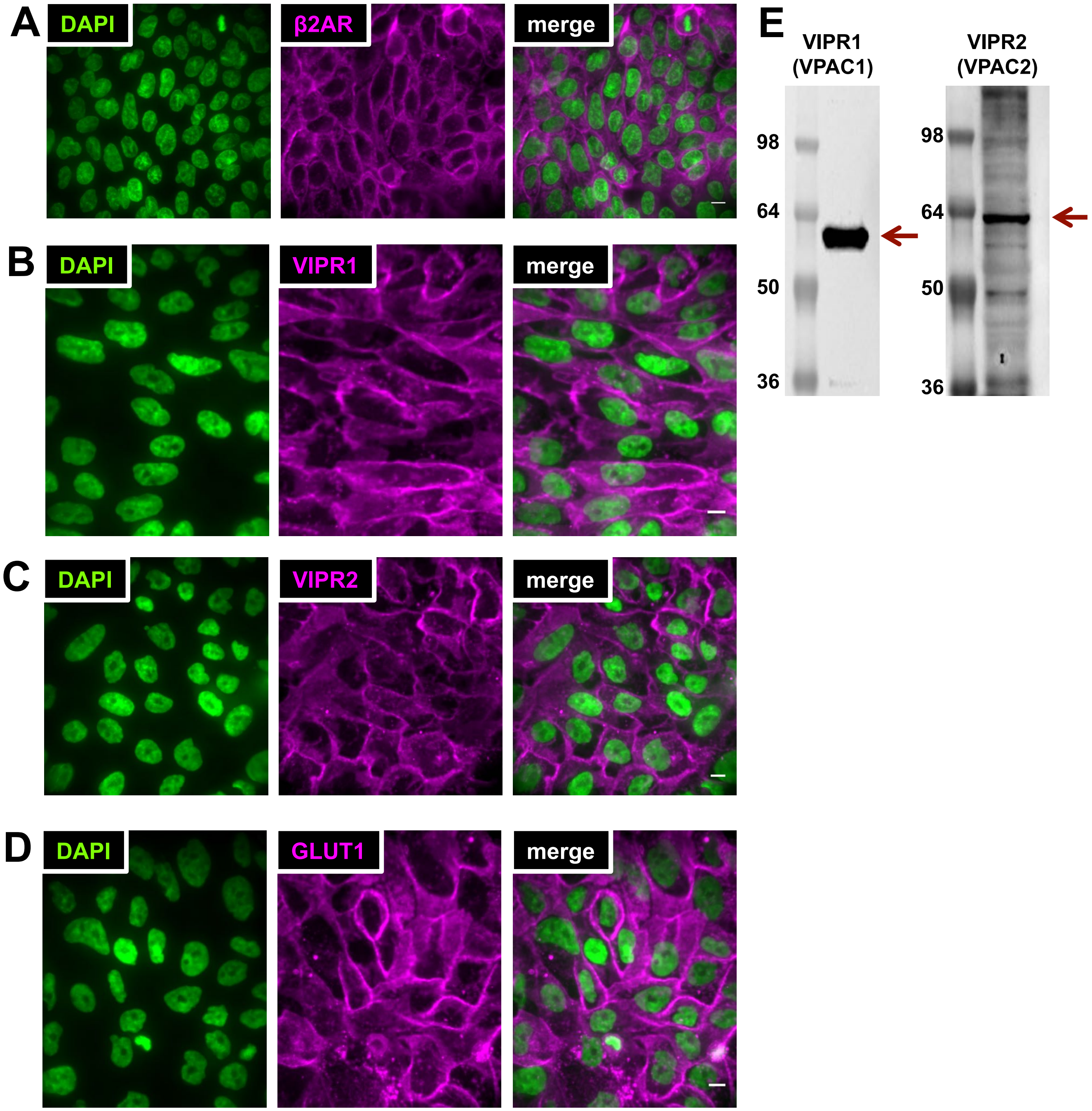


**Supplemental Figure 5: Expression of both VIPR1 (VPAC1) and VIPR2 (VPAC2) in Calu-3 serous like cells. (A-D).** Cells were seeded and grown as a monolaryer on collagen coated glass bottom dishes (MatTek), and confluent monolayers were fixed in ice cold MeOH for 3 min before immunostaining as described in the Supplemental Methods. GLUT1 and β2AR1 were used as plasma membrane controls. **(E).** Western blot showing bands corresponding to VIPR1 and VIPR2 using antibodies from *B* and *C* and as used it the main text.

#### Supplemental Figure 6

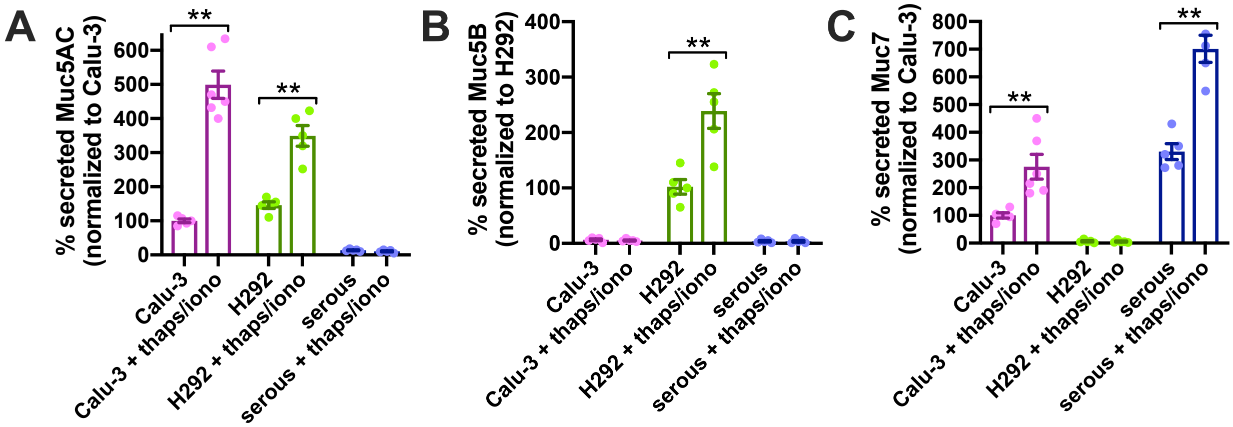


**Supplemental Figure 6: Verification of Muc7 but not Muc5AC or Muc5B production from primary human nasal serous cell cultures, suggesting maintenance of serous phenotype.** ASL was collected ± stimulation with thapsigarin and ionomycin (10 µg/ml each; 30 min, basolaterally) to maximally elevate $Ca^{2+}$ and activate acute secretion. **(A).** Calu-3 cells produced goblet cell Muc5AC (47) as previously reported (48-50), as did H292 cells, as previously reported (51-54). **(B).** H292 cells produced mucous cell-marker Muc5B (55, 56), as previously reported (57-60). Calu-3 and serous cells did not make detectible Muc5B. **(C).** Both Calu-3 and primary serous cells produced serous cell marker Muc7 (55, 56). H292 cells did not. Secretion of all mucins was increased acutely after basolateral stimulation with thapsigargin and ionomycin. All data are mean ± SEM of 3-5 independent experiments using primary serious from 3-5 separate non-CF patients.

#### Supplemental Figure 7

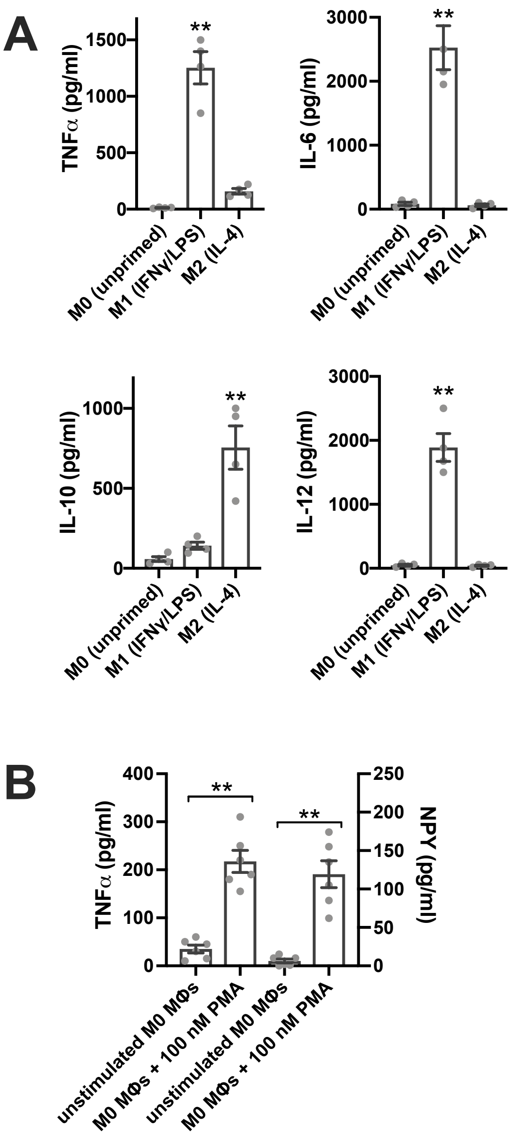


**Supplemental Figure 7: Confirmation of macrophage (MΦ) differentiation by production of appropriate cytokines in response to M1 vs M2 polarization. (A).** Human monocyte-derived MΦ were cultured as described in the text, and stimulated as indicated in the graphs for the final 3 days of the 10 day differentiation. M1 polarization (IFN γ + LPS (61, 62)) resulted in robust secretion of TNFα, IL-6, and IL-12, while M2 polarization (IL-4 (61, 62)) resulted in robust secretion of IL-10, as determined by ELISA. **(B).** Stimulation of M0 (unpolarized) MΦs by PMA for 48 hrs. resulted in secretion of TNFα (left two bars) as well as NPY (right two bars), as previously reported (63-67), as determined by ELISA. Add data are from 6 independent experiments from MΦs isolated from 3 separate individuals (2 experiments per individual). Significance determined by 1-way ANOVA with Bonferroni posttest; ***p* <0.01.

#### Supplemental Figure 8

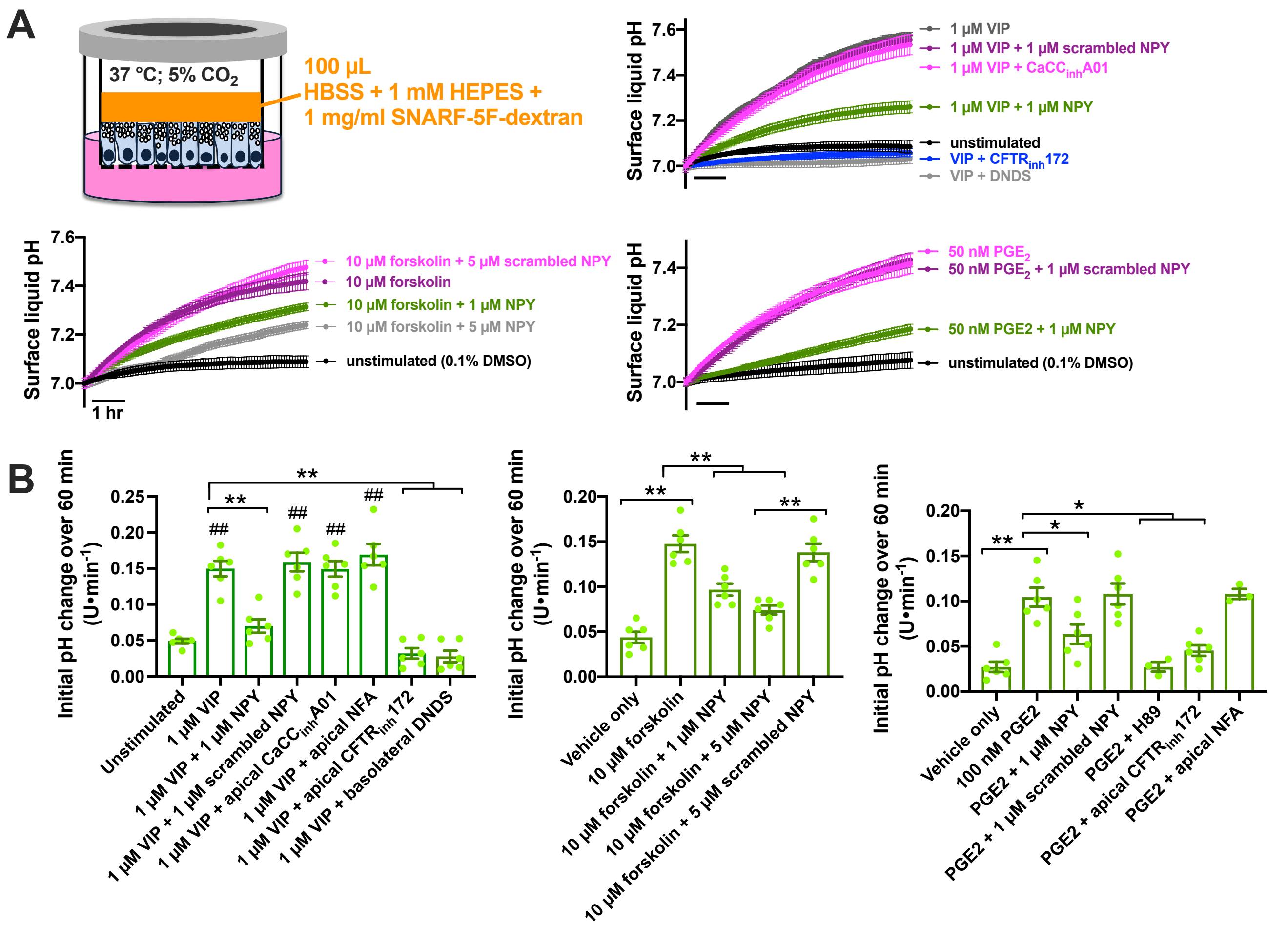


**Supplemental Figure 8. VIP, forskolin, or prostaglandin $E_2$ ($PGE_2$) increased apical secretion of $HCO_3^-$ over >8 hours, while NPY reduced secretion in response to all three agonists. (A).** Non-CF serous cell ALIs were imaged in a stage top incubator (Tokai Hit) ta 37 °C with 5% $CO_2$. 100 uL of 1 mg/ml SNARF dextran in low buffering capacity solution was added (HBSS with 1 mM HEPES, as described (22)), and pH was measured every 10 min. These experiments facilitated addition of inhibitors to the apical side. Shown are representative traces (average of 3 ALIs, 3 fields per ALI) from single experiments. (B). Bar graphs (mean ± SEM) of 6 individual experiments from at least 3 patients. Left bar graph shows inhibition of VIP-induced $pH_i$ increase by NPY but not scrambled NPY as well as by $CFTR_{inh}172$ (15 µM) but not by apical TMEM16A inhibitors $CaCC_{inh}$-A01 (15 µm) or niflumic acid (NFA; 100 µM). Block by basolateral DNDS (25 µM) suggests surface liquid $pH_i$ increases are due to $HCO_3^-$ secretion sustained by $Na^+HCO_3^-$ co-transporter (NBC) activity. Middle bar graph shows reduction of forskolin-induced ASL $pH_i$ increase by NPY but not scrambed NPY. Right bar graph shows inhibition of $PGE_2$-inducd $HCO_3^-$ secretion by NPY, protein kinase A inhibitor H89 (10 µM), or $CFTR_{inh}172$. Significance determined by 1-way ANOVA with Bonferroni posttest; ** *p*<0.01 vs bracketed bars and ## *p*<0.01 vs unstimulated control. These data support the hypothesis that cAMP-elevating agonists like VIP, forskolin, and PGE2 activates $HCO_3^-$ secretion through apical CFTR sustained by basolateral NBC and support results from the main text that NPY has an inhibitory effect on this process.

#### Supplemental Figure 9

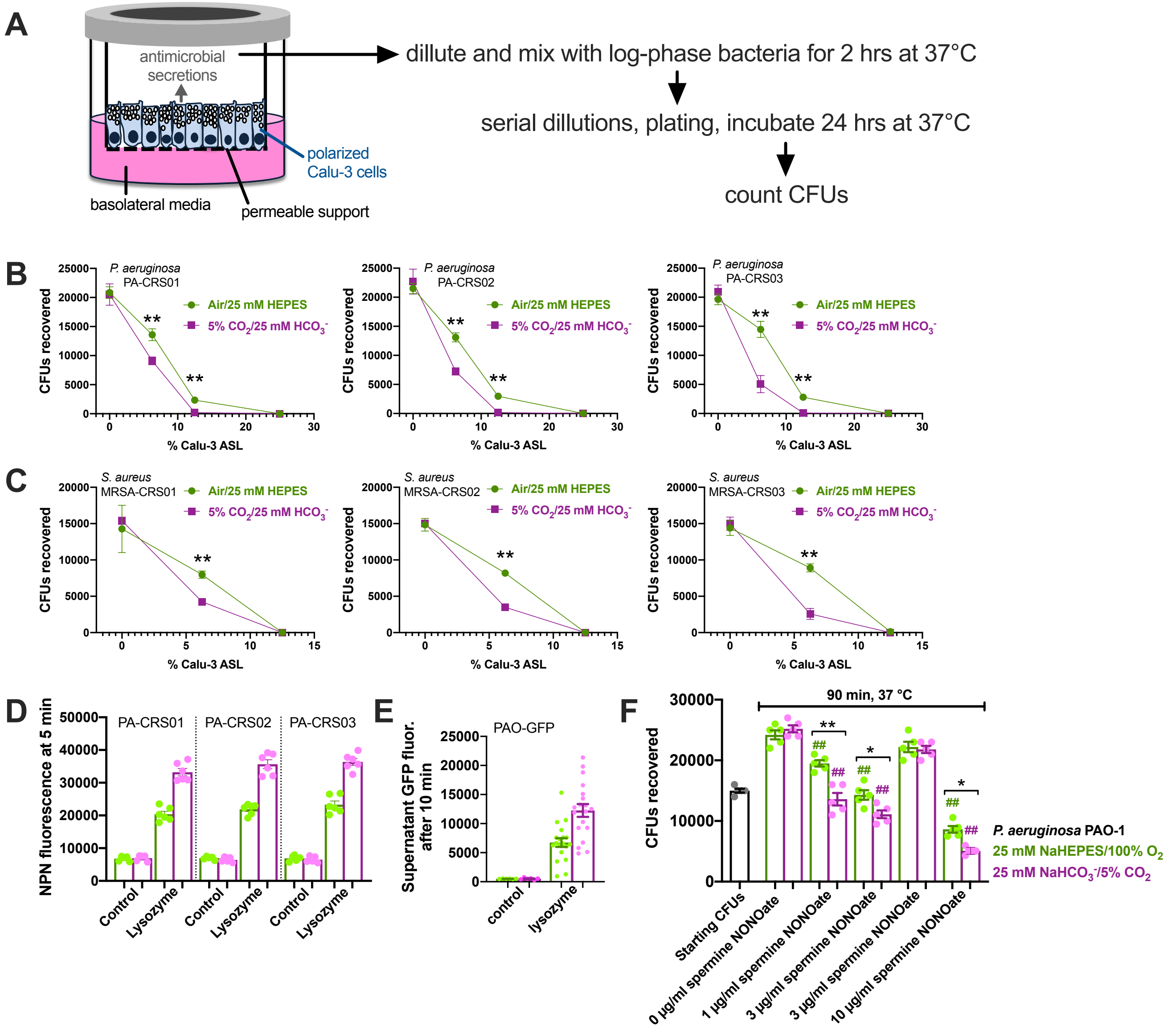


**Supplemental Figure 9. $HCO_3^-$ increases antimicrobial efficacy of Calu-3 secretions against clinical *Pseudomonas aeruginosa* strains isolated from chronic rhinosinusitis (CRS) patients.** Antimicrobial assays were carried out as described ((24) modified from(30)) using ASL from Calu-3 bronchial cell cultures. **(A-C).** Apical washings (collected with either 25% PBS + 20 mM HEPES or 25% PBS + 25 mM $HCO_3^-$) were mixed with clinical CRS isolates of *P. aeruginosa* (B) or methicillin-resistant *S. aureus* (C) and incubated at 37°C in room air (HEPES-buffered washings) or 5% $CO_2$ (HCO3$^-$ -buffered washings) followed by serial dilutions and spotting on plates for CFU counting. At low dilutions (6.25-12.5%), antimicrobial activity was greater in the presence of 5% $CO_2$. **(D).** NPN fluorescence (reflecting uptake due to cell wall damage) of clinical *P. aeruginosa* strains was measured after 5 min of lysozyme treatment (as described (24)) in the presence (pink) or absence of $HCO_3^-$ (green). **(E).** GFP-release of GFP-expressing *P. aeruginosa* (PAO-GFP) was measured during lysozyme treatment in the presence (pink) or absence (green) of $HCO_3^-$. **(F).** *P. aeruginosa* lab type strain PAO-1 was mixed in the presence (pink) or absence (green) of $HCO_3^-$ with various concentrations of NO donors, which have anti-bacterial effects (as described (68)). Overall, the presence of $HCO_3^-$ had small but significant pro-bactericidal effect in all assays tested.

#### Supplemental Figure 10

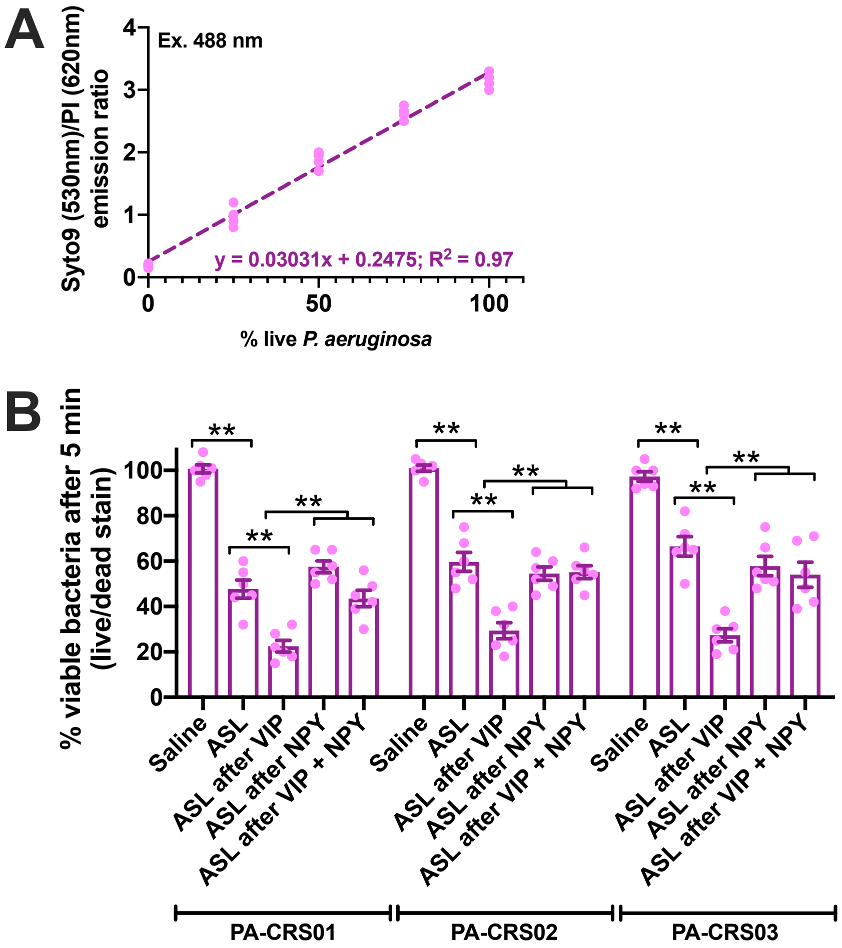


**Supplemental Figure 10: Confirmation of neuropeptide-induced changes in ASL antimicrobial efficacy by live-dead staining.** Incubation of dilutions of live and heat-killed P. aeruginosa (PAO-1) showed a linear relationship of Syto9 (live cell stain) and propidium iodide (PI; dead cell stain). Strain PA-CRS01 was used for calibration. **(B).** Live bacteria were mixed with ASL washings from primary serious ALI cultures stimulated as indicated, stained with Syto9 and PI, and read on a plate reader using 488 nm excitation and ratiometric emission (530 and 620 nm). Calibration from **A** was used to convert fluorescence ratio into viability. Bar graph shows mean ± SEM from 5-6 independent experiments using ALI washings from at least 3 separate patients. Three clinical isolates of *P. aeruginosa* isolated from CRS patients were used. Significance determined by 1-way ANOVA with Bonferroni posttest; ** p <0.01.

#### Supplemental Figure 11

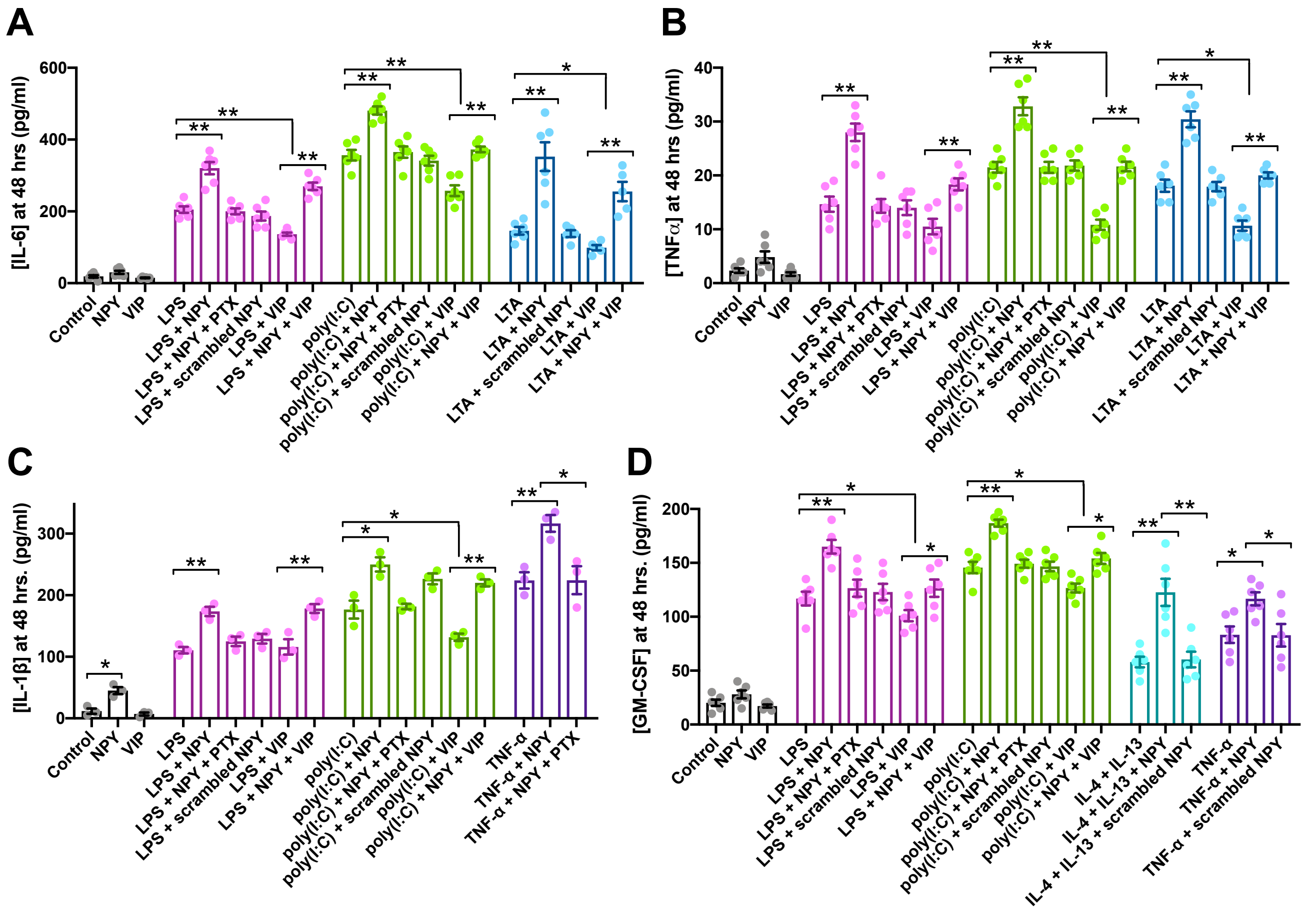


**Supplemental Figure 11: Pro-inflammatory effects of NPY and anti-inflammatory effects fo VIP.** As described in the text, serous cell cultures were stimulated with TLR4 agonist LPS (1 μg/ml), TLR3 agonist poly(I:C) (5 μg/ml), TLR2 agonist LTA (1 μg/ml), TNFα (100 ng/ml) or a Th2 cocktail of IL-4 and IL-13 (20 ng/ml each) on the apical side only, with VIP (1 μM) and/or NPY (100 nM) or scrambled NPY (100 nM) on the basolateral side, or followed by collection of basolateral media and determination of IL-6 (A), TNFα (B), IL-1β (C), or GM-CSF (D) concentration by ELISA. In most cases, NPY potentiated inflammatory responses while VIP reduce them. The only cytokine affected by either VIP or NPY alone was IL-1β, which was increased by NPY. Bar graphs show individual experiments using at least 6 ALI cultures from at least 3 patients (2 ALIs per patient); Significance determined by 1-way ANOVA with Bonferroni posttest comparing secretion of each specific cytokine among bars within each color-matched stimulus group (LPS, poly(I:C), LTA, TNFα, or IL-4 and IL-13; *p<0.05 and **p<0.01.

#### Supplemental Figure 12:

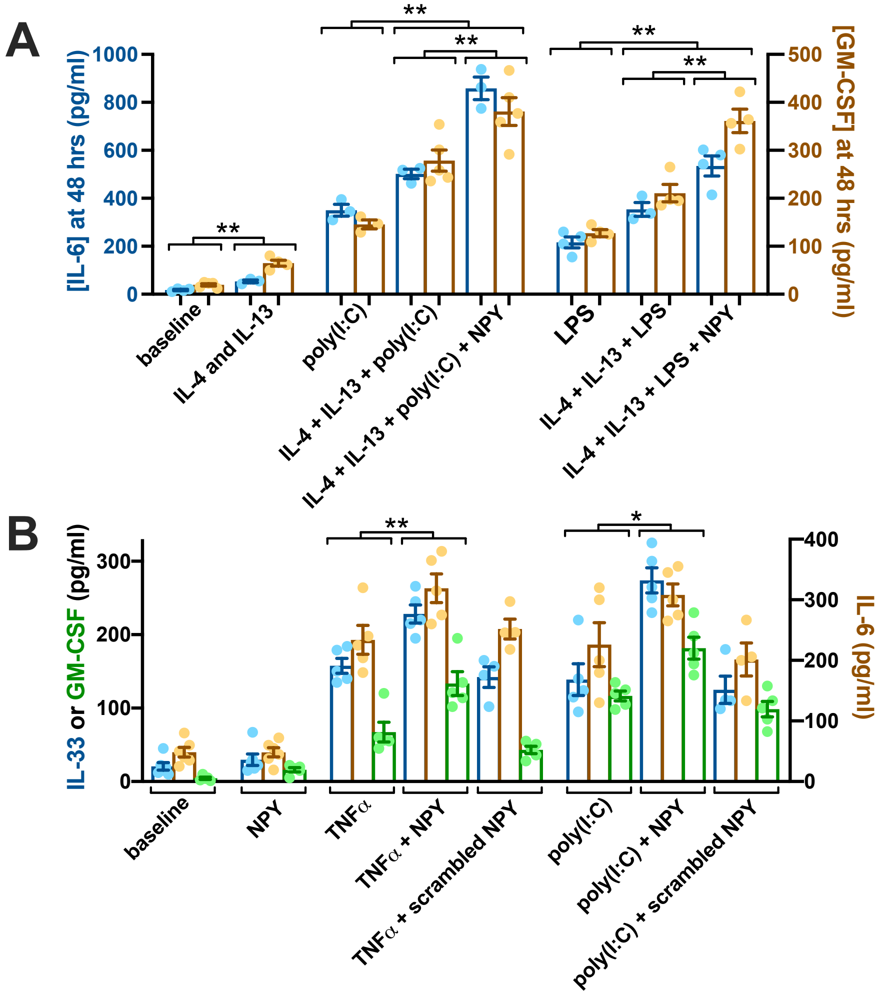


**Supplemental Figure 12: Pro-inflammatory effects of NPY in cultured (A) and acutely isolated (B) airway gland serous cells. (A).** IL-6 and GM-CSF secretion was measured by ELISA using non-CF primary serous cell ALIs stimulated with a Th2 cytokine cocktail (IL-4 + IL-13) as well as TLR3 agonists poly(I:C) or TLR4 agonist (LPS) ± NPY for 48 hrs. IL-4 + IL-13 increased responses to poly(I:C) and LPS, and NPY had a further pro-inflammatory effect even in the presence of IL-4 and IL-13, suggesting that NPY can augment inflammatory responses even in the strong Th2 environment that accompanies many inflammatory airway diseases. Bar graphs show mean ± SEM from 3-5 individual experiments each using an ALI from a separate non-CF patients. Significance determined by 1-way ANOVA with Bonferroni posttest; * $p<0.05$ and ** $p<0.01$. **(B).** Isolated acinar cells were stimulated with TNFα or poly(I:C) ± NPY or scrambled NPY. NPY, but not scrambled NPY, increased secretion of IL-33, GM-CSF, and IL-6 (measured via ELISA) in response to both TNFα or poly(I:C) but had minimal effect on its own, supporting data from cultured cells. Bar graphs show mean ± SEM from 6 individual experiments using 2 ALIs each from 3 non-CF patients. Significance determined by 1-way ANOVA with Bonferroni posttest; * $p<0.05$ and ** $p<0.01$.

**Supplemental Figure 13:**

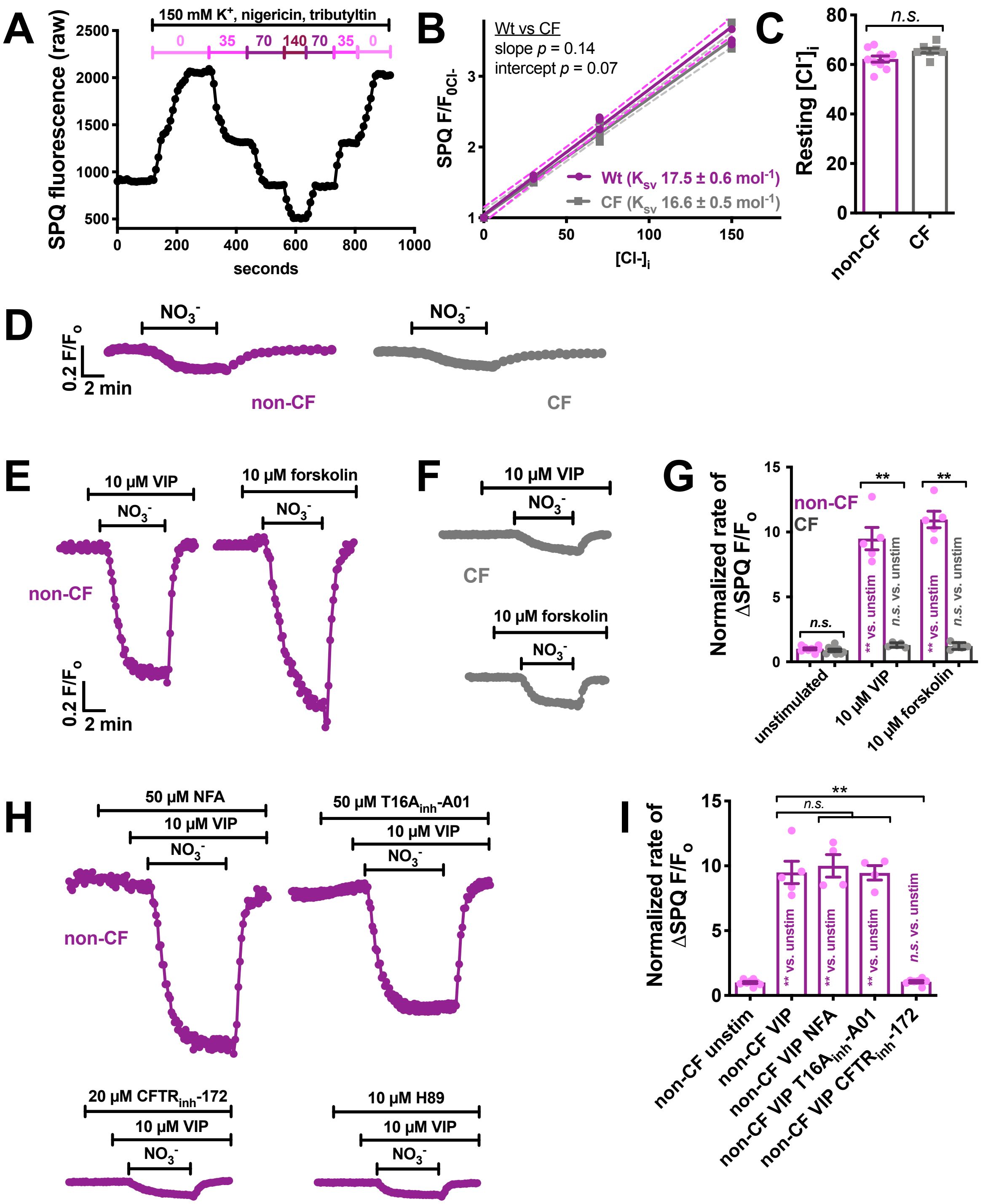

**Supplemental Figure 13: Resting $[Cl^-]_i$ is not different in non-CF and CF serous cells, but CF serous cells lack VIP/cAMP-stimulated $Cl^-$ permeability.** Non-CF and CF serous cells were obtained and isolated from patient samples and loaded with $Cl^-$ -sensitive dye SPQ as described in the Supplemental Methods and (1, 6, 7). **(A).** Representative trace of calibration of SPQ fluorescence at various $[Cl^-]_i$ values was carried out using high extracellular $K^+$ solution and $H^+/K^+$ exchanger nigericin and anion exchanger tributyltin. **(B).** Stern-Volmer plot (as described (1)) showed Stern-Volmer constant ($K_{SV}$) values of ~17/mol for both genotypes and revealed similar resting $[Cl^-]_i$ **(C).** Bar graph of resting $[Cl^-]_i$ (mean ± SEM) in non-CF and CF serous cells, which not significantly different by Student's *t* test. **(D-G).** We examined $Cl^-$ permeability using extracellular $NO_3^-$ substitution with SPQ loaded cells. SPQ is quenched by $Cl^-$ but not by $NO_3^-$, and $Cl^-$ channels are nearly equally permeable to $Cl^-$ and $NO_3^-$. $NO_3^-$ substitution (0-$Cl^-_o$) revealed identical resting $Cl^-$ permeabilities in non-CF and CF cells (*D, G*). However, when stimulated with VIP or forskolin, $Cl^-$ permeability increased in non-CF but not CF cells (*E-G*). A downward deflection of traces reflects a decrease in $[Cl^-]_i$ (increase in SPQ $F/F_o$). Bar graph in G shows mean ± SEM; * and ** = p< 0.05 and 0.01, respectively (One-way ANOVA with Bonferroni posttest. All data points are independent experiments from 3-4 CF and 3-5 non CF patients (at least 2 independent acinar cell experiments per patient). These data show that cAMP-activated $Cl^-$ permeability is absent in CF serous cells. **(H-I).** In non-CF cells, increased $Cl^-$ permeability in response to VIP was inhibited by $CFTR_{inh}172$ (CFTR inhibitor) or H89 (PKA inhibitor) but not by niflumic acid or $T16A_{inh}$-A01 ($Ca^{2+}$-activated $Cl^-$ channel inhibitors). Bar graph in I shows mean ± SEM; * and ** = p< 0.05 and 0.01, respectively by one-way ANOVA with Bonferroni posttest. Thus, VIP-activated $Cl^-$ permeability is cAMP-dependent and is blocked by CFTR inhibition.

#### Supplemental Figure 14

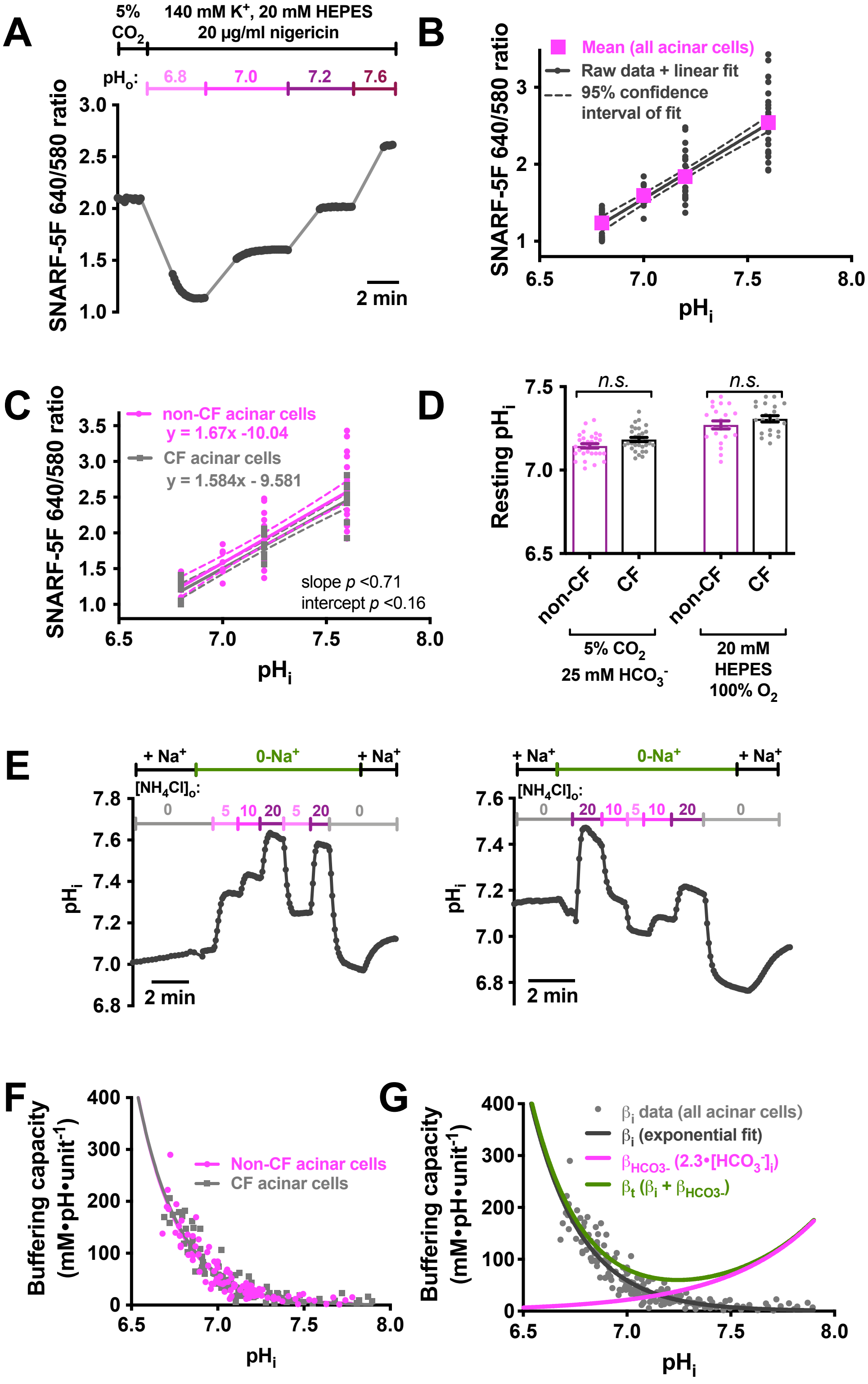

**Supplemental Figure 14: Measurement of intracellular pH ($pH_i$) in serous cells from CF and non-CF patients. (A-C).** Non-CF and CF serous cells were loaded with the ratiometric intracellular pH ($pH_i$) indicator SNARF-5F. SNARF-5F fluorescence was calibrated using high $K^+$ solutions of known pH containing $H^+/K^+$ exchanger nigericin (as described in (2)). Example calibration shown in (A). *B* shows calibration of acinar cells of all genotypes and *C* shows CF vs non-CF cells (C). No differences were observed between CF and non-CF acinar cells (n = 3 patients each), allowing comparison of the SNARF responses in the two groups. **(D).** Resting $pH_i$ was extrapolated from experiments in the presence or absence of $HCO_3^-$. No significant difference was observed in CF vs non-CF cells. Mean ± SEM; *n.s.* = not significantly different by one-way ANOVA with Bonferroni posttest. All data points are independent experiments from 3-4 CF and 3-5 non CF patients (at least 2 independent acinar cell experiments per patient). **(E-G)** Intrinsic ($HCO_3^-$-independent) $pH_i$ buffering capacity ($\beta_i$) was measured using $NH_3/NH_4$ pulse method [described in (2, 13)) under 0-$Na^+$ conditions to reduce $pH_i$ regulatory mechanisms. Because of marked variation in buffering capacity of various cell types due to size and organelle composition, $\beta_i$ must be experimentally determined. Representative calibration experiments shown in (*E*). Pooled buffering capacity measurements were used to compare CF and non-CF acinar cells (*F*). No significant difference in $\beta_i$ was observed between the two cells. This means $pH_i$ changes similarly represent $OH^-$eq fluxes in the two groups; $pH_i$ changes can thus be compared between the two groups. $\beta_i$ was fit with an exponential decay curve (G) and combined with $HCO_3^-$-dependent buffering ($\beta_{HCO3-}$) to calculate total buffering capacity ($\beta_t$) to convert $pH_i$ changes to $OH^-$ eq fluxes (not shown here). We measured pHi changes in cells exposed to solutions of various $[NH_4Cl]_o$. Exposure of cells to a solution of $NH_3$-$NH_4^+$ leads to rapid entry of membrane-permeant $NH_3$ into the cell, causing $pH_i$ alkalinization as $H^+$ is consumed as intracellular $NH_3$ converts to $NH_4^+$. This is followed by a slower acidification, likely $NH_4^+$ entry through $K^+$ channels or the $Na^+/K^+$ ATPase (12, 69). Upon changing $[NH_3]_o$, the $[NH_4^+]_i$ can be calculated using Henderson-Hasselbach with $[NH_4^+]_i = [NH_3]_i \times 10^{pKa-pHi}$ with pKa of $NH_3/NH_4^+ = 9.2$ (13). Solutions containing 0, 5, 10, and 20 mM $[NH_4Cl]_o$ contained 0, 0.6, 1.2, and 2.5 mM $[NH_3]_o$, respectively, at $pH_o = 7.4$. Buffering was calculated after an experimental change in $[NH_3]_o$ using the initial fast $pH_i$ increase or decrease to estimate buffering power around the midpoint of the $pH_i$ change $\Delta[NH_4^+]/\Delta pH_i$ (units of mmol•$L^{-1}$ of acid or base equivalent required to change $pH_i$ by one unit). Raw data points (*E*) were fit with an exponential decay function. No overt difference was observed between CF and non-CF acinar cells in $\beta_i$. Total buffering capacity ($\beta_t$; *G*) was calculated using all data points (both genotypes) and adding the $\beta_i$ curve to $\beta_{HCO3-}$ (2.3 x $[HCO_3^-]$, with $[HCO_3^-]_i$ calculated from Henderson Hasselbach with $CO_2$ clamped at 5%).

#### Supplemental Figure 15

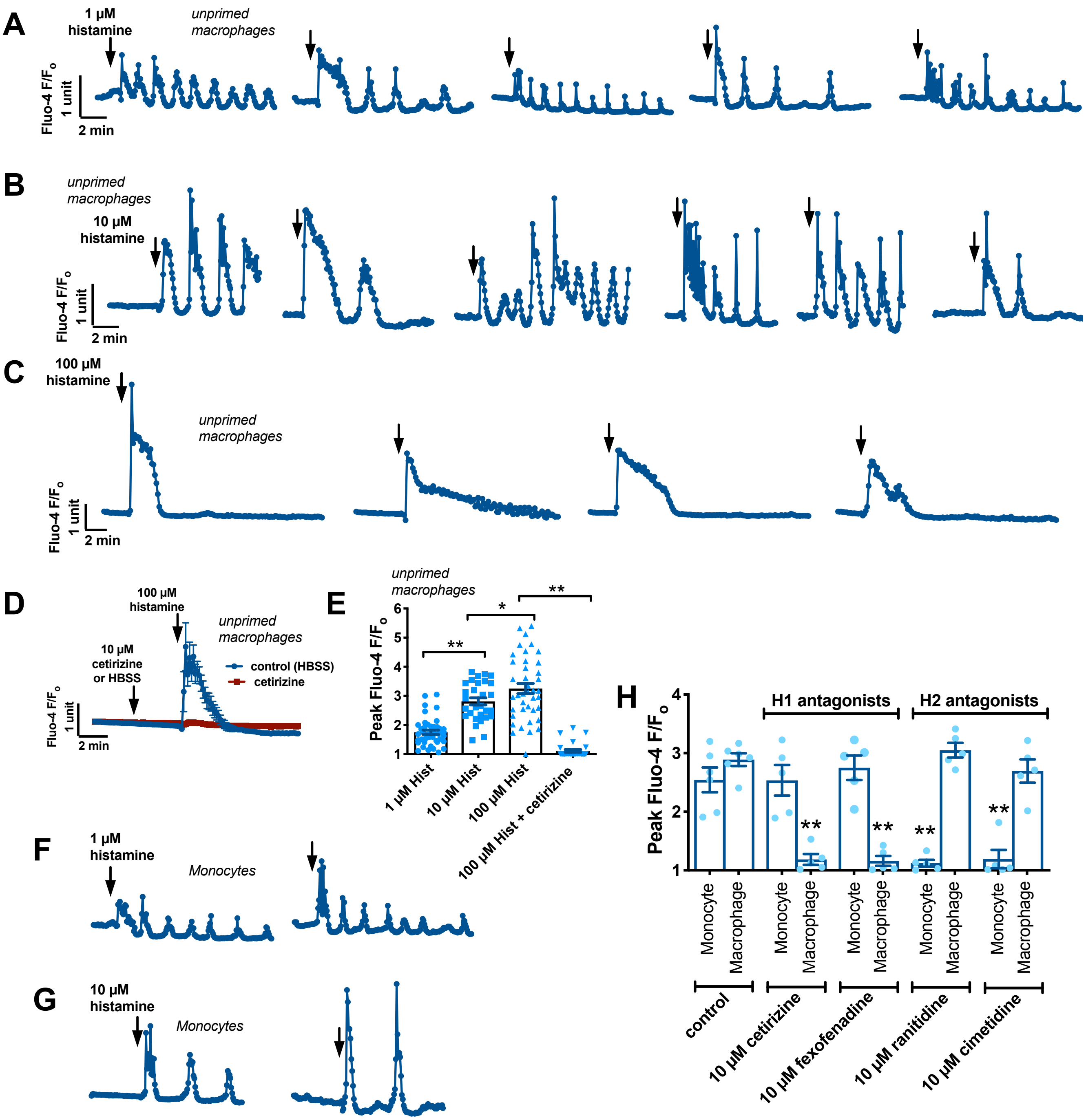

**Supplemental Figure 15: Confirmation of MΦ differentiation by functional H1 receptor expression.** Differentiation of monocytes into MΦs is accompanied by switch of histamine receptor expression from H2 to H1 isoform (70-72). **(A-C).** Representative $Ca^{2+}$ oscillations induced in individual Fluo-4 loaded MΦs by 1, 10 µM histamine as well as larger transients with 100 µM histamine in MΦs differentiated for 10 days as indicated in the text. **(D)** Average representative traces (~25 MΦs) of response to 100 µM histamine in the absence (blue) or presence of 10 µM cetirizine (H1 antagonist). (E) Plot of responses from individual MΦs from at least 3 independent experiments using MΦs from at least 3 individuals. **(F-G).** Representative $Ca^{2+}$ oscillations from freshly isolated monocytes imaged on Cell-Tak-coated coverslips. **(H).** Bar graph of individual experiments (n = 3-6 from at least 3 patients) showing inhibition of $Ca^{2+}$ responses to histamine by H1 antagonists cetirizine or fexofenadine in MΦs and H2 antagonists ranitidine and toitidine in monocytes. Bar graphs are mean ± SEM with significance determined by 1-way ANOVA with Bonferroni posttest; * $p<0.05$ and ** $p<0.01$.
